# TRACE-Seq Reveals Clonal Reconstitution Dynamics of Gene Targeted Human Hematopoietic Stem Cells

**DOI:** 10.1101/2020.05.25.115329

**Authors:** Rajiv Sharma, Daniel P Dever, Ciaran M Lee, Armon Azizi, Yidan Pan, Joab Camarena, Thomas Köhnke, Gang Bao, Matthew H Porteus, Ravindra Majeti

## Abstract

Targeted DNA correction of disease-causing mutations in hematopoietic stem and progenitor cells (HSPCs) may usher in a new class of medicines to treat genetic diseases of the blood and immune system. With state-of-the-art methodologies, it is now possible to correct disease-causing mutations at high frequencies in HSPCs by combining ribonucleoprotein (RNP) delivery of Cas9 and chemically modified sgRNAs with homologous DNA donors via recombinant adeno-associated viral vector serotype six (AAV6). However, because of the precise nucleotide-resolution nature of gene correction, these current approaches do not allow for clonal tracking of gene targeted HSPCs. Here, we describe <u>T</u>racking <u>R</u>ecombination <u>A</u>lleles in <u>C</u>lonal <u>E</u>ngraftment using <u>seq</u>uencing (TRACE-Seq), a novel methodology that utilizes barcoded AAV6 donor template libraries, carrying either in-frame silent mutations or semi-randomized nucleotide sequences outside the coding region, to track the *in vivo* lineage contribution of gene targeted HSPC clones. By targeting the *HBB* gene with an AAV6 donor template library consisting of ∼20,000 possible unique exon 1 in-frame silent mutations, we track the hematopoietic reconstitution of *HBB* targeted myeloid-skewed, lymphoid-skewed, and balanced multi-lineage repopulating human HSPC clones in immunodeficient mice. We anticipate that this methodology has the potential to be used for HSPC clonal tracking of Cas9 RNP and AAV6-mediated gene targeting outcomes in translational and basic research settings.

## Introduction

Genetic diseases of the blood and immune system, including the hemoglobinopathies and primary immunodeficiencies, affect millions of people worldwide with limited treatment options. Clinical development of *ex vivo* lentiviral (LV)-mediated gene addition in hematopoietic stem and progenitor cells (HSPCs) has demonstrated that a patient’s own HSPCs can be modified and re-transplanted to restore proper cell function in the hematopoietic system^1^. While no severe adverse events have been reported resulting from insertional mutagenesis in more than 200 patients transplanted with LV *ex vivo* manipulated HSPCs^2^, efficacy in restoring protein/cell function and ultimately disease amelioration has varied. In some diseases, this lack of therapeutic efficacy is possibly the result of irregular spatiotemporal transgene expression due to the semi-random integration patterns of LVs.

Tracking the transgene integration sites (IS) by deep sequencing has been used to “barcode” clones in heterogeneous cell populations that contribute to blood reconstitution in the human transplantation setting. In clinical trials, IS methodology has been used to track genetically modified memory T-cells^3^, waves of hematopoietic repopulation kinetics^4^, as well as dynamics and outputs of HSPC subpopulations in autologous graft composition^5^. These seminal studies provided new insights into the reconstitution of human hematopoiesis following autologous transplantation. Importantly, IS can also provide evidence of potential concerning integration patterns in tumor-suppressor genes, like *PTEN*^*6*^, TET2^7^ and *NF1*^*8*^, which can be closely monitored during long-term follow-up to predict future severe adverse events.

Genetic barcoding on the DNA level has been used to track the *in vitro*^*9*^ and *in vivo*^*10-13*^ clonal dynamics of heterogeneous mammalian cellular populations and offers several advantages over lentiviral IS tracking, although it has not been used clinically. First, the amplified region is known and nearly the same for each barcode simplifying recovery from targeted cells, as opposed to semi-random LV integrations, which require amplification of unknown sequences. Second, it is far less likely for differences in amplification efficiency or secondary structure to lead to drop off or mis-quantification of clone sizes^14^. Altogether, genetic barcoding, combined with high-throughput sequencing, can enable sensitive and quantitative assessment of heterogeneous cell populations.

Genome editing provides an alternative approach to lentiviral integrations to perform permanent genetic engineering of cells. Genome editing can be performed using non-nuclease approaches^15, 16^, by base editing^17^, or by prime-editing^18^, but the most developed and efficient form of precision engineering in human cells utilizes engineered nuclease-based approaches^19-24^. The repurposing of the bacterial CRISPR/Cas9 system for use in human cells^25, 26^ has democratized the field of genome editing because of its ease of use, high activity, and high specificity, especially using high fidelity versions of Cas9^27^. Nuclease-based editing has now entered clinical trials with more on the horizon^28^.

Genome editing by combining ribonucleoprotein (RNP, Cas9 protein complexed to synthetic stabilized, single guide RNAs) combined with the use of the non-integrating AAV6 viral vector to deliver the donor template has been shown to be a highly effective system to modify therapeutically relevant primary human cells including HSPCs, T-cells, and induced pluripotent cells^29^. This approach has shown pre-clinical promise to usher in a new class of medicines for sickle cell disease^27, 30^, SCID-X1^31, 32^, MPS I^33^, chronic granulomatous disease^34^, X-linked Hyper IgM^35^, and cancer^36^. The specificity of genome editing, however, means that with current approaches it is not possible to track the output of any specific gene modified cell. The spectrum of non-homologous end joining (NHEJ)-introduced INDELs is also not broad enough to reliably measure clonal dynamics within a population^37^. Yet, understanding clonal dynamics within large populations of engineered cells is important and significant in both pre-clinical studies and potentially clinical studies. Therefore, we developed a barcode system for homologous recombination-based genome editing. We applied this system to understand the clonal dynamics of CD34^+^ human HSPCs following transplantation into immunodeficient NSG mice.

We describe TRACE-Seq, a methodology that allows for both correction of disease-specific mutations and for the tracking of contributions of gene targeted HSPCs to single and multi-lineage hematopoietic reconstitution. In brief, we demonstrate: 1) design and production of barcoded AAV6 donor templates using silent in-frame mutations or semi-randomized nucleotides outside the coding region (but inside the homology arms), 2) barcoding the first 9 amino acids of HBB exon 1 with ∼20,000 possible AAV6 donor templates maintains high gene correction frequencies while preserving robust beta globin expression levels, 3) the ability to track the reconstitution of gene corrected myeloid- and lymphoid-skewed HSPC clones as well as balanced multi-lineage clones, and 4) an analysis pipeline that includes a highly adaptable platform for interpreting and summarizing rich datasets from clonal tracking studies that is deployable as a website accessible to researchers with no coding experience. TRACE-Seq demonstrates that Cas9 RNP and AAV6-mediated gene correction can be used to target a single HSC clone that can then robustly repopulate the myeloid and lymphoid branches of the hematopoietic system. This method and information further supports the translational potential of homologous recombination based approaches for the treatment of genetic diseases of the blood and immune system.

## Methods

### Donor design and cloning

#### HBB barcode donor libraries

AAV transfer plasmid with inverted terminal repeats (ITR) from AAV2 that contained 2.4kb of the *HBB* gene previously described^30^ was digested with NcoI and BamH1 restriction enzymes (NEB) that resulted in deletion of a 435bp band and the digested backbone was collected for further subcloning. Double stranded DNA gBlock (IDT) pools with degenerate bases representing silent mutations containing 645 bases of homology were ordered in four separate oligo pools (as detailed below with bold depicting silent mutation region). Four different barcoded dsDNA oligo pools were ordered to maximize potential silent mutations that if all were ordered in the same library would have resulted in amino acid changes to the coding region. Each HBB barcoded dsDNA pool was then digested with NcoI and BamHI resulting in a 435bp band that was collected and purified. NEB Assembly ligation reactions were performed for 1 hour at 50°C using digested, gel purified vector. Ligated HBB barcoded donor pools were transformed using NEB DH10B electrocompetent bacteria (NEB C3020K) or XL10-Gold competent cells (Agilent 200315) according to the manufacturer’s protocol. At least two times the theoretical maximum number of possible barcoded donor templates were plated to ensure generation of as much diversity as possible. Endotoxin-free maxipreps were generated for AAV6 production and purification. As noted, *HBB* barcode pool 3 was not included in genome editing experiments because enrichment of the original undigested donor plasmid was seen during sequencing of the plasmid library.

HBB barcode pool 1 (8192 possible unique donor templates):

agaagagccaaggacaggtacggctgtcatcacttagacctcaccctgtggagccacaccctagggttggccaatctactcccaggagcagggagggcaggagccagggctgggcataaaagtcagggcagagccatctattgcttacatttgcttctgacacaactgtgttcactagcaacctcaa acagacaccatgg**<u>TNCAYTTRACNCCNGARGARAARTCNGCAGTCACT</u>**gccctgtggggcaaggtgaa cgtggatgaagttggtggtgaggccctgggcaggttggtatcaaggttacaagacaggtttaaggagaccaatagaaactgggcatgtggagacagagaagactcttgggtttctgataggcactgactctctctgcctattggtctattttcccacccttaggctgctggtggtctacccttggacccagaggttctttgagtcctttggggatctgtccactcctgatgctgttatgggcaaccctaaggtgaaggctcatggcaagaaagtgctcggtgcctttagtgatggcctggctcacctggacaacctcaagggcacctttgccacactgagtgagctgcactgtgacaagctgcacgtggatcctgagaacttca gggtga

HBB barcode pool 2 (4096 possible unique donor templates):

agaagagccaaggacaggtacggctgtcatcacttagacctcaccctgtggagccacaccctagggttggccaatctactcccaggagcagggagggcaggagccagggctgggcataaaagtcagggcagagccatctattgcttacatttgcttctgacacaactgtgttcactagcaacctcaa acagacaccatgg**<u>TNCAYTTRACNCCNGARGARAARAGYGCAGTCACT</u>**gccctgtggggcaaggtgaa cgtggatgaagttggtggtgaggccctgggcaggttggtatcaaggttacaagacaggtttaaggagaccaatagaaactgggcatgtggagacagagaagactcttgggtttctgataggcactgactctctctgcctattggtctattttcccacccttaggctgctggtggtctacccttggacccagaggttctttgagtcctttggggatctgtccactcctgatgctgttatgggcaaccctaaggtgaaggctcatggcaagaaagtgctcggtgcctttagtgatggcctggctcacctggacaacctcaagggcacctttgccacactgagtgagctgcactgtgacaagctgcacgtggatcctgagaacttca gggtga

HBB barcode pool 3 (16384 possible unique donor templates):

agaagagccaaggacaggtacggctgtcatcacttagacctcaccctgtggagccacaccctagggttggccaatctactcccaggagcagggagggcaggagccagggctgggcataaaagtcagggcagagccatctattgcttacatttgcttctgacacaactgtgttcactagcaacctcaa acagacaccatgg**<u>TNCAYCTNACNCCNGARGARAARTCNGCAGTCACT</u>**gccctgtggggcaaggtgaa cgtggatgaagttggtggtgaggccctgggcaggttggtatcaaggttacaagacaggtttaaggagaccaatagaaactgggcatgtggagacagagaagactcttgggtttctgataggcactgactctctctgcctattggtctattttcccacccttaggctgctggtggtctacccttggacccagaggttctttgagtcctttggggatctgtccactcctgatgctgttatgggcaaccctaaggtgaaggctcatggcaagaaagtgctcggtgcctttagtgatggcctggctcacctggacaacctcaagggcacctttgccacactgagtgagctgcactgtgacaagctgcacgtggatcctgagaacttca gggtga

HBB barcode 4 (8192 possible unique donor templates):

agaagagccaaggacaggtacggctgtcatcacttagacctcaccctgtggagccacaccctagggttggccaatctactcccaggagcagggagggcaggagccagggctgggcataaaagtcagggcagagccatctattgcttacatttgcttctgacacaactgtgttcactagcaacctcaa acagacaccatgg**<u>TNCAYCTNACNCCNGARGARAARAGYGCAGTCACT</u>**gccctgtggggcaaggtgaa cgtggatgaagttggtggtgaggccctgggcaggttggtatcaaggttacaagacaggtttaaggagaccaatagaaactgggcatgtggagacagagaagactcttgggtttctgataggcactgactctctctgcctattggtctattttcccacccttaggctgctggtggtctacccttggacccagaggttctttgagtcctttggggatctgtccactcctgatgctgttatgggcaaccctaaggtgaaggctcatggcaagaaagtgctcggtgcctttagtgatggcctggctcacctggacaacctcaagggcacctttgccacactgagtgagctgcactgtgacaagctgcacgtggatcctgagaacttca gggtga

#### AAVS1 barcode donor libraries

AAVS1 barcode libraries were generated similarly to HBB libraries. Briefly, degenerate nucleotides (following the pattern “VHDBVHDBVHDB,” in order to minimize homopolymer stretches as described^38^ were introduced by PCR 3’ of the mTagBFP2 reporter cassette. pAAV-MCS plasmid (Agilent Technologies) containing ITRs from AAV serotype 2 (AAV2) was digested with NotI and barcode-containing PCR fragments were assembled into the backbone using NEB Assembly using the following primers, prior to transformation with XL10-Gold competent cells (Agilent 200315):

Insert_Fw1: CCATCACTAGGGGTTCCTGCGGCCGCCACCGTTTTTCT

Insert_Rv1: TTAATTAAGCTTGTGCCCCAGTTTGCTAGG

Insert_Fw2: TGGGGCACAAGCTTAATTAA**<u>VHDBVHDBVHDB</u>**CTCGAGGGCGC

Insert_Rv2: CCATCACTAGGGGTTCCTGCGGCCGCAGAACTCAGGAC

### AAV6 production and purification

HBB barcoded recombinant adeno-associated virus serotype (AAV6) six vectors were produced and purified as previously described^39^. Briefly, 293FT cells (Life Technologies) were seeded at 15 million cells per dish in a total of ten 15-cm dishes one to two days before transfection (or until they are 80-90% confluent). One 15-cm dish was transfected with 6μg ITR-containing *HBB* barcoded donor plasmid pools 1-4 and 22μg pDGM6. Cells were incubated for 48-72h until collection of AAV6 from cells by three freezes-thaw cycles. AAV6 vectors were purified on an iodixanol density gradient, AAV6 vectors were extracted at the 60-40% iodixanol interface, and dialyzed in PBS with 5% sorbitol with 10K MWCO Slide-A-Lyzer G2 Dialysis Cassette (Thermo Fisher Scientific). Finally, vectors were added to pluronic acid to a final concentration of 0.001%, aliquoted, and stored at −80°C until use. AAV6 vectors were tittered using digital droplet PCR to measure the number of vector genomes as described previously^40^. AAVS1 barcoded AAV6 donors were produced as described above but purified using a commercial purification kit (Takara Bio #6666).

### CD34^+^ hematopoietic stem and progenitor cell culture

All CD34^+^ cells used in these experiments were cultured as previously described^39^. In brief, cells were cultured in low-density conditions (<250,000 cells/mL), low oxygen conditions (5% O_2_), in SFEMII (Stemcell Technologies) or SCGM (CellGenix) base media supplemented with 100ng/mL of TPO, SCF, FLT3L, IL-6 and the small molecule UM-171 (35nM). For *in vitro* studies presented in **Figure 2**, CD34^+^ cells from sickle cell disease patients were obtained as a kind gift from Dr. John Tisdale at the National Institute of Health (that were mobilized with plerixafor in accordance with their informed consent) or from routine non-mobilized peripheral blood transfusions at Stanford University under informed consent. For *in vivo* studies presented, cord blood-derived CD34^+^ cells were purchased from AllCells or Stemcell Technologies and were thawed according to the manufacturer’s recommendations.

**Figure 1.**
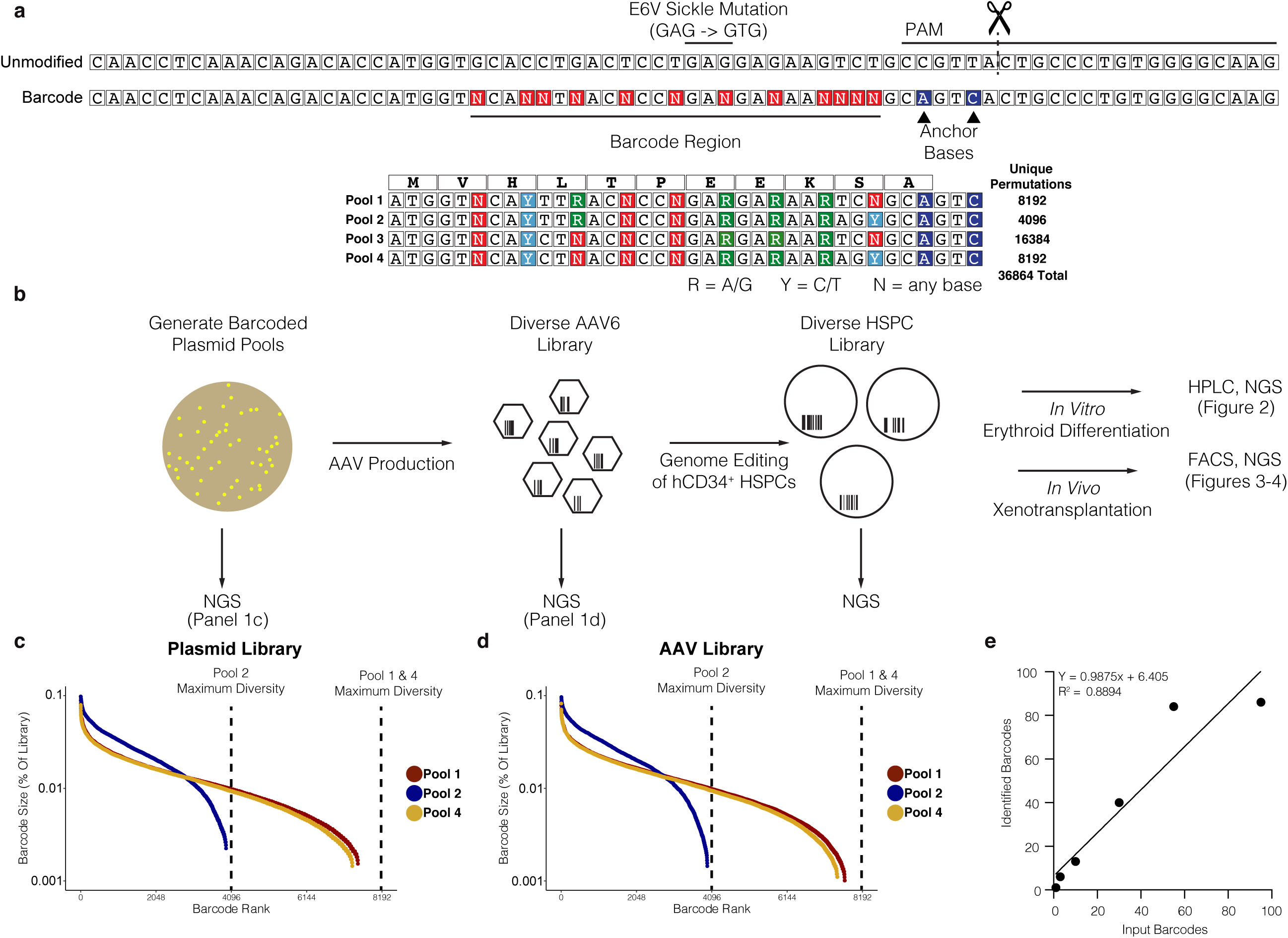
Design and production of barcoded AAV6 donors for long-term genetic tracking of gene targeted cells and their progeny. **a** Schematic of HBB targeting strategy. Top: Unmodified (WT) and barcoded HBB alleles depicted, with location of the E6V (GAG -> GTG) sickle cell disease mutation and CRISPR/Cas9 target sites labeled. Bottom: β-globin ORF translation with four barcode pools representing all possible silent mutations encoding amino acids 1-9. **b** Schematic of barcode library generation and experimental design. **c/d** Percentages of reads from each valid barcode identified through amplicon sequencing of plasmids (c) and AAV (d) pools 1, 2, and 4. **e** Recovery of barcodes from untreated genomic DNA containing 1, 3, 10, 30, and 95 individual plasmids containing HBB barcodes. Expected number of barcodes are plotted against the number of barcodes called by the TRACE-seq pipeline after filtering.

**Figure 2.**
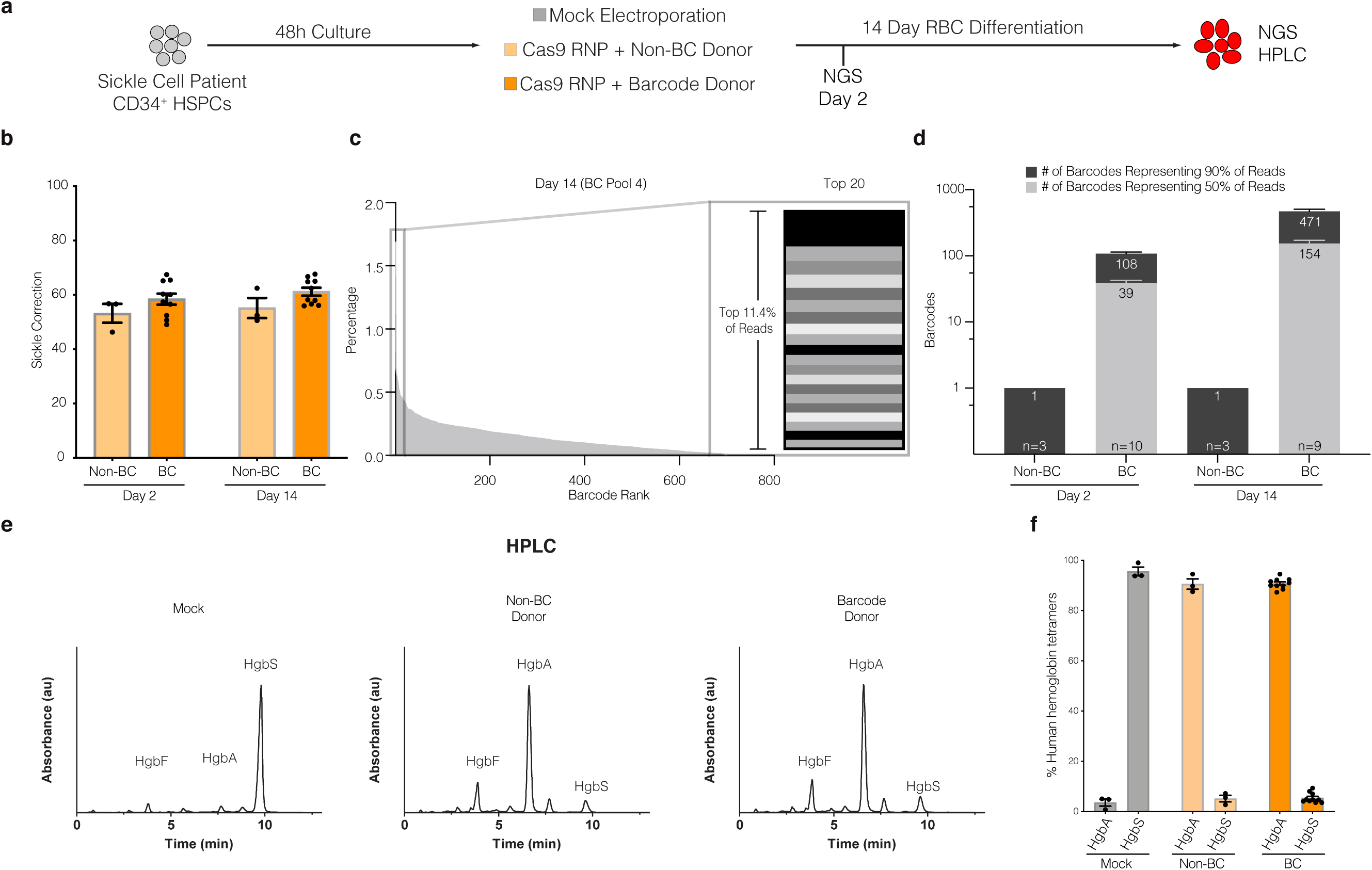
*Correction of the Sickle Cell Disease-causing E6V mutation using barcoded AAV6 donors in SCD-derived* CD34^+^ HSPCs. **a** Experimental design – SCD patient derived CD34^+^ HSPCs edited with CRISPR/Cas9 RNP and electroporation only (mock), single donor (non-BC), or barcode donor (BC) AAV6 HDR templates. **b** SCD correction efficiency (percentage of corrected sickle cell alleles) of non-BC and BC treated groups as a fraction of total NGS reads (e.g. HR reads / [sum of HR reads + unmodified reads].) **c** Representative example of barcode fractions in descending order from one donor at day 14 time point. Right: Top 20 clones represented as stacked bar graph (representing 11.4% of reads). **d** Number of unique barcode alleles comprising the top 50% and top 90% of reads from each treatment condition, sampling approximately 1000 cells per condition (see **Supplemental Table 1**). **e** Representative hemoglobin tetramer HPLC chromatograms of RBC differentiated cell lysates at day 14 post treatment. **f** Quantification of total hemoglobin protein expression in each group. Each data point represents an individual biological replicate. HgbA: adult hemoglobin HgF: fetal hemoglobin HbS: sickle hemoglobin. AAV6: Recombinant AAV2/6 vector.

### Cas9/sgRNA and AAV6-mediated genome editing

All experiments in these studies used the R691A HiFi Cas9 mutant^27^ (IDT and Aldevron), and chemically synthesized guide RNA (sgRNA)^41^ (Synthego). The guide sequences were as follows: HBB: 5′-CTTGCCCCACAGGGCAGTAA-3′ and AAVS1: 5′-GGGGCCACTAGGGACAGGAT-3′. Genome editing experiments using Cas9/sgRNA and AAV6 were performed as previously described^39^. In brief, CD34^+^ HSPCs were thawed and plated for 48h to allow for recovery of freezing process and pre-stimulation of cell cycle. CD34^+^ HSPCs were then electroporated in 100μl electroporation reaction buffer P3 (Lonza) with 30μg HiFi Cas9 and 16μg MS sgRNA (pre-complexed for 10 minutes at room temperature; HiFi RNP). HSPCs were resuspended with HiFi RNP in P3 buffer and electroporated using program DZ-100 on the Lonza 4D nucleofector. Immediately following electroporation, CD34^+^ HSPCs were transduced with HBB-specific AAV6 barcoded donor template libraries at 2500-5000 vector genomes per cell and 20000 vector genomes per cell for AAVS1-specific AAV6 barcoded libraries. 12-16h post transduction, targeted cells were washed and resuspended in fresh media and allowed to culture for additional 24-36h, with a total manufacturing time less than 96h.

### In vitro erythrocyte differentiation of HBB-targeted CD34^+^ HSPCs

SCD-HSPCs were targeted with either the therapeutic AAV6 donor (with one sequence) or the HBB barcoded AAV6 donor template library and subjected to the *in vitro* erythrocyte differentiation protocol two days post targeting as previously described^27, 42, 43^. Base medium was supplemented with 100U/mL of penicillin–streptomycin, 10ng/mL SCF, 1ng/mL IL-3 (PeproTech), 3U/mL erythropoietin (eBiosciences), 200μg/mL transferrin (Sigma-Aldrich), 3% antibody serum (heat-inactivated from Atlanta Biologicals, Flowery Branch, GA, USA), 2% human plasma (umbilical cord blood), 10μg/mL insulin (Sigma Aldrich) and 3U/mL heparin (Sigma-Aldrich). Briefly, targeted HSPCs were differentiated into erythrocytes using a three-phase differentiation protocol that lasted 14-16 days in culture. The first phase of erythroid differentiation corresponded to days 0-7 (day 0 being day 2 after electroporation). During the second phase of differentiation, corresponding to days 7-10, IL-3 was discontinued from culture medium. In the third and final phase, corresponding to days 10-16, transferrin was increased to 1mg/mL. Differentiated cells were then harvested for analysis of hemoglobin tetramers by cation-exchange high performance liquid chromatography.

### Hemoglobin tetramer analysis via cation-exchange HPLC

Hemoglobin tetramer analysis was performed as previously described^27^. Briefly, red blood cell pellets were flash frozen post differentiation until tetramer analysis where pellets were then thawed, lysed with 3 times volume of water, incubated for 15 minutes and then sonicated for 30 seconds to finalize the lysing procedure. Cells were then centrifuged for 5 minutes at 13,000 rpm and used for input to analyze steady-state hemoglobin tetramer levels. Transfused blood from sickle cell disease patients was always used to ascertain the retention time of sickle, adult and fetal human hemoglobin.

### Transplantation of targeted CD34^+^ HSPCs into NSG mice

Six to eight week old immunodeficient NSG female mice were sublethally irradiated with 200cGy 12-24h before injection of cells. For primary transplants, 2-4 x 10^5^ targeted CD34^+^ HSPCs were harvested two days post electroporation, spun down at 300g, and resuspended in 25μl PBS before intrafemoral transplantation into the right femur of female NSG mice. For secondary transplants, mononuclear cells (MNCs) were harvested from primary transplanted NSG mice, and half of the total MNCs were used to transplant one sublethally irradiated female NSG mouse via tail vein injection.

### Analysis of human engraftment and fluorescent activated cell sorting

16-18 weeks following transplantation of targeted HSPCs, mice were euthanized, bones (2x femurs, 2x pelvis, 2x tibia, sternum, spine) were collected and crushed as previously described^30, 39^. MNCs were harvested by ficoll gradient centrifugation and human hematopoietic cells were identified by flow cytometry using the following antibody cocktail: HLA-A/B/C FITC (clone W6/32, Biolegend), mouse CD45.1 PE-CY7 (clone A20, Thermo Scientific), CD34 APC (clone 581, Biolegend), CD33 V450 (clone WM53, BD Biosciences), CD19 Percp5.5 (clone HIB19, BD Biosciences), CD10 APC-Cy7 (HI10a, Biolegend), mTer119 PeCy5 (clone Ter-119, Thermo Scientific), and CD235a PE (HIR2, Thermo Scientific). For mice transplanted with AAVS1-edited HSPCs, the following cocktail was used: HLA-A/B/C FITC (clone W6/32, Biolegend), mouse CD45.1 PE-CY7 (clone A20, Thermo Scientific), CD34 APC (clone 581, Biolegend), CD33 PE (clone WM53, BD Biosciences), CD19 BB700 (clone HIB19, BD Biosciences), CD3 APC-Cy7 (clone SK7, BD Biosciences). For AAVS1-edited HSPCs, CD33^Hi^ and CD33^Mid^ were sorted individually, however the data were aggregated for analysis. Human hematopoietic cells were identified as HLA-A/B/C positive and mCD45.1 negative. The following gating scheme was used to sort cell lineages to be analyzed for barcoded recombination alleles: Myeloid cells (CD33^+^), B Cells (CD19^+^), HSPCs (CD10^−^, CD34^+^, CD19^-^, CD33^-^), and erythrocytes (Ter119^-^, mCD45.1^-^, CD19^-^, CD33-, CD10^−^, CD235a^+^). Sorted cells were spun down, genomic DNA was harvested using QuickExtract (Lucigen), and was saved until library preparation and sequencing.

### Sequencing library preparation

Harvested cells were lysed using QuickExtract DNA Extraction Solution (Lucigen, Cat. No. QE09050) following manufacturers protocol. Based on the starting cell count, 0.5-1μL QuickExtract lysate was used for PCR. All PCRs for library preparation were carried out using Q5 High-Fidelity 2X Master Mix (NEB, Cat. No. M0492L). An initial enrichment amplification of 15 cycles was followed with a second round of PCR using unique P5 and P7 indexing primer combinations for 15 cycles and purified using 1.8X SPRI beads. For nested PCR, an initial amplification of 30 cycles was used. PCR products were analyzed by gel electrophoresis and purified using 1X SPRI beads.

PCR products were normalized, pooled and then gel extracted using the QIAEX II Gel Extraction Kit (Qiagen, Cat. No. 20051). The resulting libraries were sequenced using both Illumina Miseq (2 x 150 bp paired end) and Illumina HiSeq 4000 (2 x 150 bp paired end) platforms. Illumina HiSeq 4000 sequencing were performed by Novogene Corporation.

### Index switching correction of false positive NGS reads

We utilized two independent methods to determine the incidence of index-switching present in samples that were run on a HiSeq 4000^44, 45^. In one approach, we calculated the number of contaminating reads between two different amplicons sequenced in the same pool. As a second approach, we utilized the algorithm developed by Larrson et al. to estimate the fraction of reads which were spread to other samples through index switching^46^. Both of these methods yielded an index switching incidence of 0.3%. We performed a conservative correction for this by subtracting 0.3% x [# Barcode Reads] from each barcode in each sample. We performed this correction after clustering as described in Extended Data Figure 1.

### Statistical analysis

All statistical tests used in this study were performed using GraphPad Prism 7/8 or R version 3.6.1. For comparing the average of two means, we used the Student’s t-test to reject the null hypothesis (P < 0.05).

## Results

### Design, production, and validation of barcoded AAV6 donor templates for targeting the HBB gene in human HSPCs

We previously developed an *HBB* AAV6 homologous donor template that corrects the sickle cell disease-causing mutation in HSPCs with high efficiencies^30^. Using this AAV6 donor as a template, we designed an *HBB* barcoded AAV6 donor library with the ability to: 1) correct the E6V sickle mutation, 2) preserve the reading frame of the beta globin gene, and 3) generate enough sequence diversity to track cellular events on the clonal level (throughout the manuscript we will consider unique barcodes representative of cellular clones, with the caveat that clone counts may be overestimated due to bi-allelic targeting of two barcodes into the genome of a single cell). We designed the donor pool to contain mixed nucleotides that encode silent mutations within the first 9 amino acids of the HBB coding sequence (“VHLTPEEKS”, **Figure 1a**). Using this strategy, we designed double stranded DNA oligos that contained the library of nucleotide sequences and cloned four separate pools of donors with a theoretical maximum number of 36,864 in-frame, synonymous mutations (**Figure 1b**).

To ensure that the initial plasmid library reached the theoretical maximum diversity with near-equal representation of all sequences, we performed amplicon sequencing on the initial plasmid pools. Sequencing of HBB barcoded pools 1, 2, and 4 (**Figure 1a, bottom**) revealed a wide distribution of sequences with no evidence of any highly overrepresented barcodes (**Figure 1c**). Barcode pool 3 was eliminated for further study, because it was contaminated with uncut vector control and therefore skewed barcode diversity. After validating that the plasmid pools were diverse and lacked enrichment of any one sequence, we used the HBB barcoded library plasmid pools 1, 2, and 4 to produce libraries of AAV6 homologous donor templates. After generating barcoded AAV6 donor libraries, we performed amplicon-based NGS to determine the diversity and distribution of sequences. Similar patterns were observed, suggesting standard AAV6 production protocols do not introduce donor template bias in the barcoded pool (**Figure 1d**).

### Establishing thresholds for HBB barcode quantification

Understanding the clonal dynamics of hematopoietic reconstitution through sequencing requires the ability to differentiate between low frequency barcodes and noise introduced by sequencing error. Therefore, we used a modified version of the TUBAseq pipeline to cluster cellular barcodes and differentiate between sequencing error and bona-fide barcode sequences^47^. The overall schema of the TRACE-seq pipeline is depicted in **Extended Data Figure 1a**. Briefly, we merged paired-end fastq files using the PEAR algorithm with standard parameters^48^, and then aligned reads to the human HBB gene. Reads were binned into three categories: unmodified alleles (wildtype), non-homologous end joining (NHEJ) alleles, and homologous recombination (HR) alleles. Reads were classified as unmodified if they aligned to the reference HBB gene with no genome edits. Reads were classified as NHEJ if there were any insertions or deletions within 20bp of the cut site, and if anchor bases (PAM-associated bases changed after successful HR) were unmodified (**Figure 1a**). Finally, reads were classified as HR if they had modified anchor bases and were not classified as NHEJ (**Figure 1a**). All subsequent analyses were performed exclusively on the HR reads.

To differentiate between *bona fide* barcodes and sequencing errors, variable barcode regions and non-variable training regions were extracted from the HR reads and TUBAseq was used to train an error model and cluster similar barcodes together using the DADA2 algorithm^47^. We chose a DADA2 clustering omega parameter of 10^−40^ because: 1) we found that at this omega value, the number of unfiltered barcodes called began to reach the minimum number of barcodes called per sample as omega was decreased, and 2) we found that varying this parameter did not ultimately affect the number or sequence of called barcodes after filtering (described subsequently) for samples with known barcode content (**Extended Data Figure 1b**).

In order to benchmark our analysis pipeline, we cultured individual barcoded bacterial plasmid colonies in 96 well plates and generated pooled plasmid libraries to generate a set of ground-truth samples with known barcode content. These libraries were spiked into untreated human gDNA and were subjected to our optimized amplicon sequencing and analysis pipeline. We found that clustering eliminated more than 97% of low-level noise barcodes across all samples with known barcode content, but left a small percentage of low-level barcodes in the clustered barcode set (**Extended Data Figure 1c**). Using the ground-truth samples, we determined a “high confidence” barcode threshold of 0.5%, which allowed us to quantitatively recover the expected numbers of barcodes (R^2^ = 0.89) (**Figure 1e, Extended Data Figure 1c**).

Overall, our pipeline allowed us to process raw amplicon sequencing data and generate a set of barcodes unlikely to contain spurious signals. Conceptually, we extracted barcodes from each read and eliminated barcodes which appeared to be derived from sequencing or other error using a clustering-based methodology and evidence-based filtering heuristics, resulting in a set of high-confidence barcodes with which we performed further analyses.

### Barcoding HBB exon 1 with in-frame silent mutations preserves hemoglobin expression while allowing cell tracking within a heterogeneous population

To evaluate whether the barcoded AAV6 donor libraries preserved the open reading frame of HBB following targeted integration, we compared HSPCs targeted with a non-barcoded homologous donor (containing a single corrective AAV6 genome^30^; non-BC) or a barcode donor library (BC) as illustrated in **Figure 2a**. We performed gene-targeting experiments by electroporating HiFi Cas9 and HBB-specific chemically modified guide sgRNAs^41^ into primary CD34^+^ HSPCs isolated from patients with sickle cell disease (which contained the E6V point mutation). We observed similar gene correction efficiencies between HSPCs targeted with non-BC and BC donors as quantified by amplicon-based next generation sequencing from approximately 1000 cells from each timepoint (**Figure 2b**). To assess barcode diversity, we ranked barcodes by read percentages from largest to smallest for each treatment group (**Figure 2c**). Focusing specifically on the top 20 barcodes in the representative example in **Figure 2c**, it is evident that even with a relatively small sample, we observe a fairly even distribution of barcodes, with no evidence of extreme overrepresentation from any particular sequences. We calculated the number of the most abundant barcodes comprising 50% and 90% of total HR reads as a measure of sequence diversity. As expected, the single non-BC donor sample contained one barcode (the corrected E6V sequence) along with intentional synonymous mutations^30^ that represented >94% of reads (**Figure 2d**). Of note, the remaining reads appeared to be sequencing/PCR artifacts as they often contained nonsynonymous mutations in the HBB reading frame (**Supplemental Table 1a and 1b**). In contrast, the 90^th^ percentile of barcode reads in BC donor targeted cells contained a mean of 107.7 ± 9.6 barcodes at day 2 and 471.0 ± 54.1 at day 14 (**Figure 2d**). These unique barcode counts were not surprising given the limited numbers of input cells analyzed, and the additional complexity of performing nested PCR reactions to avoid contamination from unintegrated (episomal) AAV6 donor genomes, especially at early timepoints before the cells could undergo many rounds of division. Indeed, by aggregating together all experimental replicates treated with BC donors, the 90^th^ percentile of barcode reads contained >3200 barcodes (data not shown), suggesting barcode identification was limited by sampling depth. Importantly, the barcodes observed in the BC donor treated samples preserved the HBB coding sequence even though their sequences varied greatly (**Supplemental Table 1c and 1d**). These results are consistent with the notion that targeting HSPCs with a BC donor produces a diverse pool of HSPCs capable of correcting the E6V sickle mutation, and that diversity is maintained within a two-week period of *in vitro* culture.

While the sequencing data suggest that the HSPCs targeted with the BC donors exhibit robust E6V gene correction frequencies, the introduction of silent mutations may interfere with hemoglobin protein expression. To assess this possibility, we performed *in vitro* erythroid differentiation of non-BC and BC targeted HSPCs and collected red blood cell pellets for HPLC analysis of hemoglobin tetramer formation. While the unedited mock sample contained >90% sickle hemoglobin (HgbS) (of total hemoglobin), HSPCs targeted with non-BC or BC AAV6 donors both exhibited >90% adult hemoglobin (HgbA) protein production (**Figure 2e-f**). These results suggest the silent mutations introduced by the BC donor had no significant negative influence on overall translation efficiency, despite being produced from a diverse pool of >450 unique sequences in the bulk-edited population.

### TRACE-Seq reveals long-term engraftment of lineage-specific and bi-lineage potent HBB targeted hematopoietic stem and progenitor cells

In addition to correcting the E6V mutation and restoring HgbA expression, barcoded AAV6 donors can be utilized to label and track cells in a heterogeneous pool of HSPCs. To track cellular lineages in a pool of HBB-labeled HSPCs, we transplanted BC and non-BC control targeted cord blood CD34^+^ HSPCs via intra-femoral injection into sublethally irradiated adult female NSG recipient mice (2-4 x 10^5^ cells per mouse from n=6 total cord blood donors, see **Supplemental Table 2**). Upon sacrifice (16-18 weeks post-engraftment), mice in both transplantation groups exhibited no statistically significant differences in total human engraftment (46 ± 10.4 vs. 50 ± 10.1, non-BC and BC, respectively, **Figure 3a**). Similarly, no significant differences were seen between non-BC and BC mice in terms of lineage reconstitution of the human cells engrafted, which mainly consisted of B cells (CD19^+^), myeloid cells (CD33^+^) or HSPCs (CD19^-^CD33^-^CD10^−^CD34^+^) (**Figure 3b**).

**Figure 3.**
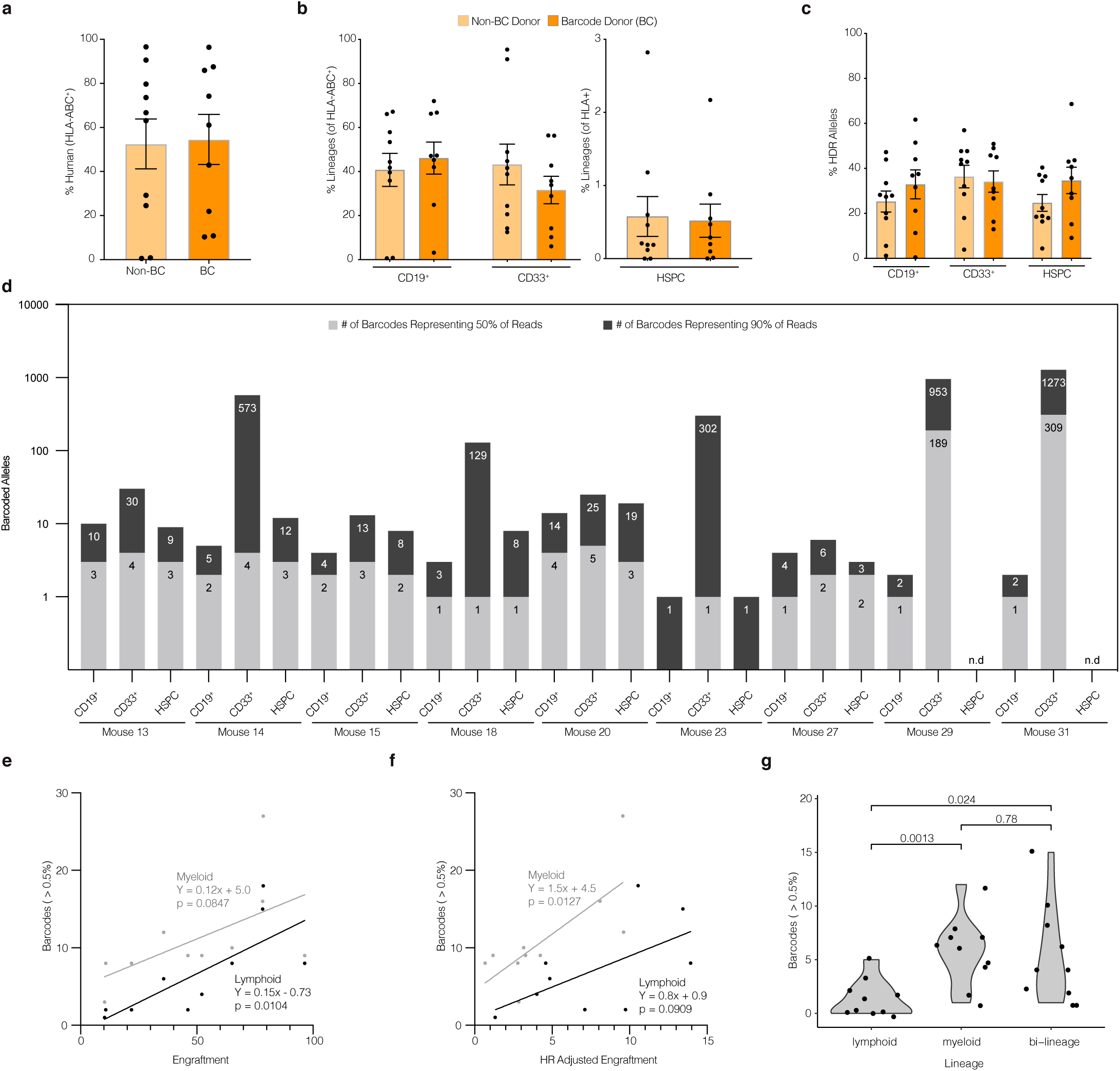
*TRACE-Seq identifies lineage-restricted and multi-potent gene targeted* HSPCs in primary NSG transplants. CD34^+^ enriched cord blood-derived HSPCs were cultured in HSPC media containing SCF, FLT3L, TPO, IL-6, and UM-171 for 48h, electroporated with Cas9 RNP (HBB sgRNA), transduced with AAV6 donors (either BC or non-BC), and cultured for an additional 48h prior to intrafemoral transplant into sublethally irradiated NSG mice (total manufacturing time was less than 96 hours). 16-18 weeks post transplantation, total BM was collected and analyzed for engraftment by flow cytometry, sorted on lineage markers, and sequenced for unique barcodes. Two independent experiments were performed to assess reproducibility of identifying clonality of gene-targeted HSPCs. **a** Total human engraftment in whole bone marrow, (as measured by proportion of human HLA-ABC^+^ cells). **b** Multilineage engraftment of human CD19^+^, CD33^+^, and HSPCs (CD19^-^CD33^-^CD10^−^CD34^+^). **c** Genome editing efficiency in each indicated sorted human lineage subset as determined by NGS (HR reads / [sum of HR reads + unmodified reads]). **d** Barcodes from each subset were sorted from largest to smallest by percentage of reads. Depicted are the numbers of most abundant, unique barcode alleles comprising the top 50% and top 90% of reads from each lineage of all mice transplanted with BC donor edited HSPCs. Mean ± SEM genomes analyzed from each group: CD19^+^: 8500 ± 1000, CD33^+^: 8800 ± 800, HSPC: 1500 ± 500 (see **Supplemental Table 2b**). **e** Correlation between numbers of high confidence barcodes (> 0.5%) in lymphoid (grey) and myeloid (black) compartments and total human engraftment (as percent of human and mouse BM-MNCs). Lymphoid and myeloid values plotted for n=9 primary engrafted mice and n=1 secondary engrafted mouse. **f** Correlation between numbers of high confidence barcodes (> 0.5%) in lymphoid (grey) and myeloid (black) compartments and HR adjusted engraftment ([human engraftment] x [lineage specific engraftment] x [HR efficiency]). Lymphoid and myeloid values plotted for n=9 primary engrafted mice and n=1 secondary engrafted mouse. **g** Numbers of high confidence barcodes from each mouse which contribute to lymphoid only (CD19^+^), myeloid only (CD33^+^), or both lineages. High confidence barcodes: barcodes with at least 0.5% representation (see **Extended Data Figure 1**). All points represent individual mice, with the exception of panels e-g (where barcodes from each mouse are separated based on lineage contribution). Error bars depict mean ± SEM. p values reflect 2-tailed t-test.

To evaluate the efficiency of non-BC or BC gene targeting in long-term engrafting HSPCs, bone marrow MNCs were sorted by flow cytometry into lineages CD19^+^ and CD33^+^, as well as the multipotent HSPC (CD19^-^CD33^-^CD10^−^CD34^+^) populations (**Extended Data Figure 2a**). Using the pipeline outlined in **Extended Figure 1**, we performed amplicon based NGS to quantify the proportions of gene targeted alleles relative to total editing events that included NHEJ and unmodified alleles. We did not detect any significant differences in the efficiency of HDR within any of these subsets between non-BC and BC donors (**Figure 3c**).

Because there was robust engraftment of HBB targeted alleles in the BC mice, we were able to track the recombination alleles within the lymphoid, myeloid, and multipotent HSPC subpopulations. We analyzed cells from a total of 9 mice sorted on lymphoid (CD19^+^), myeloid (CD33^+^), and HSPC (CD19^-^CD33^-^CD10^−^CD34^+^) markers. 130.6 ± 62.3 unique barcodes accounted for 90% of the reads with a median of 2 unique barcodes accounting for 50% or the sequencing reads from each group (**Figure 3d**). Barcodes in all three sorted populations exhibited less diversity than was observed *in vitro*, indicating that there was a reduction in clonal complexity following engraftment into mice (**Extended Data Figure 3a**). For example, the CD19^+^ compartment from Mouse 18 contained over 60 total clones passing our thresholds, with a majority of reads coming from a single barcode (**Extended Data Figure 3b**). The number of high confidence barcodes (>0.5% of reads) was correlated with total human engraftment in the lymphoid compartment and a similar trend was observed in the myeloid compartment (p=0.08) (**Figure 3e**). The same trend was observed when we correlated barcodes with lineage specific engraftment adjusted for HR frequency (**Figure 3f**). When we subdivided these more abundant barcodes into alleles that contributed to lymphoid only, myeloid only, or bi-lineage output within the mice, we observed fewer barcodes generated from lymphoid-skewed compared to myeloid-skewed or bi-lineage HSPCs (p=0.0013 and p=0.024, respectively, **Figure 3g**). These data suggest that Cas9/sgRNA and AAV6-mediated HBB gene targeting occurs in multipotent HSPCs as well as lineage-restricted HSPCs.

The gold standard for defining human long-term hematopoietic stem cell (LT-HSC) activity is to perform secondary transplants into another sublethally irradiated NSG mouse^49^. Therefore, we compared the TRACE-Seq dynamics of a primary recipient versus a secondary recipient in mouse 20 that exhibited very high engraftment (>80% human cell engraftment). While mouse 20 had a total of 17 lymphoid and 56 myeloid clones contributing to the engraftment of gene targeted HBB cells, the majority of differentiated cell output was from relatively few clones (**Figure 4a**, left panel). Four lymphoid and five myeloid lineage barcodes accounted for 50% of the reads from each population. This trend was consistent between all mice analyzed (**Extended Data Figure 3c-k**) with each mouse displaying a unique set of HBB barcodes that all maintained the coding region (**Supplemental Table 3**). Barcode reads from the same sorted cell populations from the secondary mouse transplant revealed further reductions in clonal diversity, almost to a monoclonal state, with a single clone representing 80% or more of reads in both lymphoid and myeloid lineages (**Figure 4a**, right panel, dark blue). Interestingly, the dominant clone in the secondary transplant was not the most abundant clone in the primary mouse as it only represented 10.9% of lymphoid and 16% of myeloid alleles.

**Figure 4.**
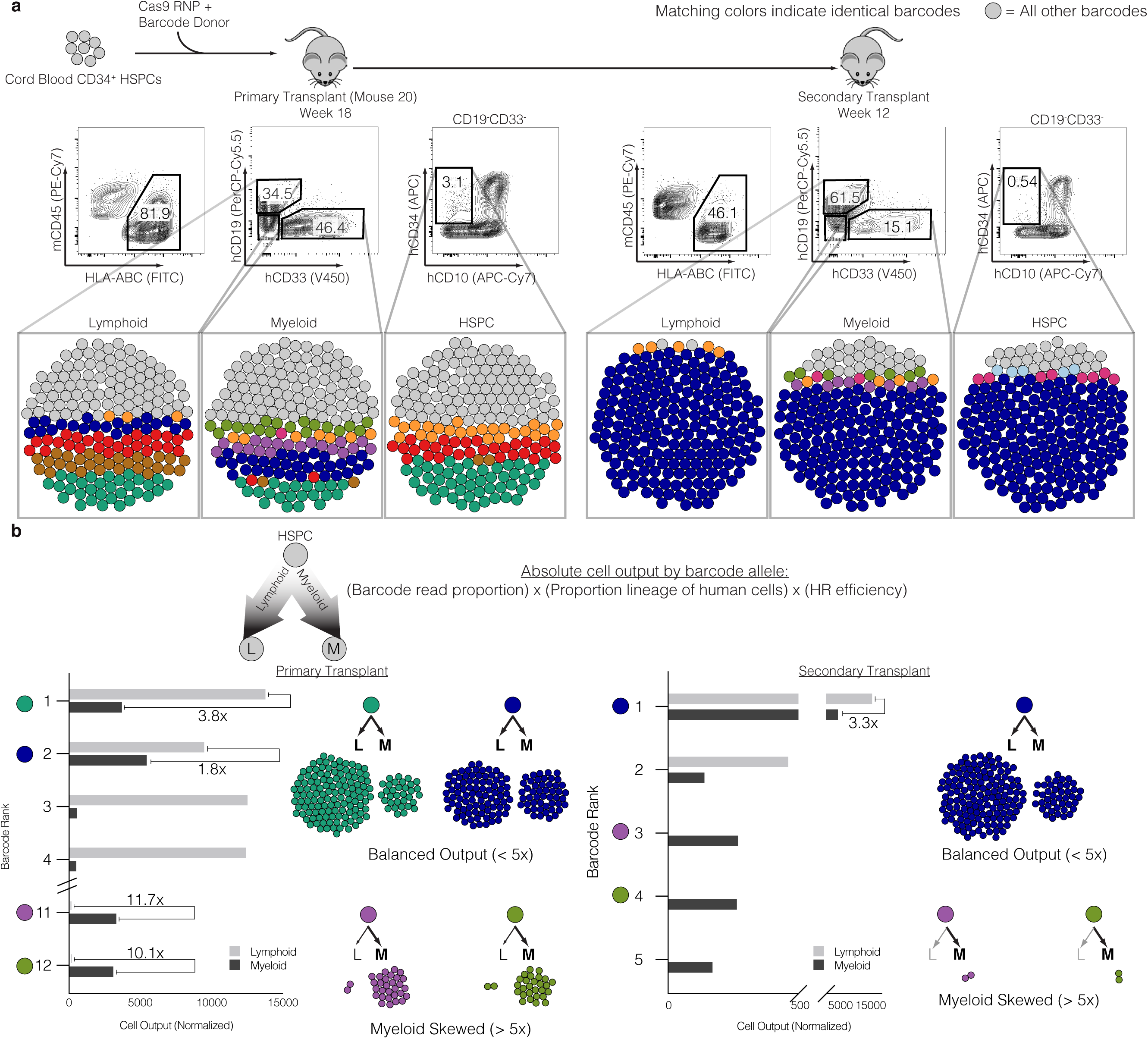
Identification of clonal dynamics of HBB-targeted HSPCs. **a** Top: Experimental schematic. Middle: Flow cytometry plots representing robust bi-lineage engraftment in primary transplant (left, week 18 post-transplant) and secondary transplant (right, week 12 post-transplant). Bottom: Bubble plots representing barcode alleles as unique colors from each indicated sorted population. Shown are the three most abundant clones from all six populations. All other barcodes represented as grey bubbles. **b** Normalized output of barcode alleles with respect to lineage contribution. Total cell output (bar graphs) from indicated barcodes adjusted for both differential lineage output and genome editing efficiency within each subset. Examples of various lineage skewing depicted, with cell counts proportional to the absolute contribution to the xenograft. Skewed output defined as 5-fold or greater bias in absolute cell counts towards lymphoid or myeloid lineages.

To understand the contribution of each clone to the absolute number of differentiated hematopoietic cells in the mouse bone marrow, we took into consideration the following parameters: 1) the fraction of unique barcode reads assigned to each clone, 2) the relative contribution of the lineage where each clone was detected to the entire graft, and 3) gene targeting frequencies (**Figure 4b**). This analysis reveals clones that are lymphoid skewed (brown and red, **Figure 4a**), myeloid skewed (purple and light green), as well as clones exhibiting balanced hematopoiesis (dark blue). We defined skewing as having a >5-fold difference in proportion between lymphoid and myeloid cells. Perhaps the most interesting observation from this analysis was that the more balanced hematopoietic clone (dark blue) was responsible for a great majority of secondary engraftment/repopulation (**Figure 4b**, right). Interestingly, while this clone contributed >80% of the engraftment of HBB targeted cells, there were still observable myeloid lineage-skewed clones present in the secondary transplant. This analysis also revealed barcode sequences that produced highly correlated read frequencies (± 2% read proportions) in both primary and secondary transplants, consistent with bi-allelic gene targeting in the same long-term HSPC (**Figure 4b**, purple and light green barcodes).

### TRACE-Seq by barcoding AAV6 donor templates outside the coding region allows for clonal tracking of AAVS1 targeted HSPCs

To test that the barcoding scheme (inside the coding region), library diversity (maximal theoretical diversity of 36,864 HBB barcodes), and/or the gene being targeted (HBB) did not bias our results, we developed a strategy to target *AAVS1* with barcoded SFFV-BFP-PolyA AAV6 donor libraries (**Extended Data Figure 4a**). We designed the AAVS1 barcoded variable region within the 3’ untranslated region of the BFP expression cassette so the barcode would be in the genomic DNA as well as mRNA. Using a design that prevents mononucleotide runs that can potentially increase sequencing error^38^, a 12 nucleotide variable barcode region resulted in a theoretical maximal barcoded AAV6 pool of 531,441 different homologous donor templates (**Extended Data Figure 4a**, bottom). Using such a large pool allowed us to rule out the possibility that the numbers of barcodes observed in the HBB system is artificially limited by the smaller diversity of the HBB barcode pool. As with the HBB pipeline (**Figure 1e**), we benchmarked our ability to differentiate sequencing error from legitimate barcodes by choosing parameters and thresholds that resulted in a high correlation between known numbers of input barcodes and barcodes identified through TRACE-seq (**Extended Data Figure 4b**).

We targeted cord blood-derived HSPCs with the AAVS1-BC pool of AAV6 donor templates and transplanted them into sublethally irradiated NSG mice to assess the clonal contribution via TRACE-Seq. Robust AAVS1-BC donor targeting into the *AAVS1* locus was achieved in two independent experiments across five HSPC donors and a mean of 2.90 ± 0.4 × 10^5^ cells transplanted per NSG mouse (**Supplemental Table 2**). Following 16-18 weeks of hematopoietic reconstitution, we observed 45.4% ± 14.2 human engraftment, with a gene targeting efficiency of 42.4% ± 11.4 (**Extended Data Figure 4c**). As with the HBB donors, the majority of differentiated cells were CD19^+^ lymphoid and CD33^+^ myeloid cells, with a strong trend towards more genome editing within the CD33^+^ population (55.8 ± 12.0 vs. 22.3 ± 11.2; p=0.06, two-tailed t-test) (**Extended Data Figure 4d, e**). To assess clonal contributions of AAVS1 targeted HSPCs, lineage specific cells (CD19^+^ or CD33^+^) were sorted (**Extended Data Figure 4e**), and AAVS1-BFP specific amplicons were generated for NGS sequencing of cells with on-target integrations of SFFV-BFP-PolyA. Consistent with our findings targeting the *HBB* locus, we identified not only similar numbers of unique barcodes (representing individual clones) in divergent hematopoietic lineages (**Extended Data Figure 4f, Figure 5a**), but also similar patterns between primary and secondary transplants, suggesting again that TRACE-Seq identifies Cas9/sgRNA and AAV6-mediated targeting of LT-HSCs (**Figure 5a, Extended Data Figure 5**). Across all mice, bi-lineage clones were seen in four out of five mice, with the exception being mouse 38, from which we were not able to sort sufficient numbers of myeloid cells for valid analysis (**Supplemental Table 4** and **Extended Data Figure 5**). As with HBB TRACE-Seq, calculating the relative cell output of individual barcodes revealed lymphoid skewed, myeloid skewed and balanced HSPC clones (**Figure 5b**, left). The most dominant clone (red), which displayed high proliferative output with a more balanced hematopoietic lineage distribution in the primary mouse, was the predominant clone in the secondary transplant (**Figure 5b**, right). In addition, we observed less abundant, myeloid skewed clones (blue and green) in both primary and secondary transplants. These results confirm that gene targeted LT-HSC clones contribute to robust multi-lineage engraftment.

**Figure 5.**
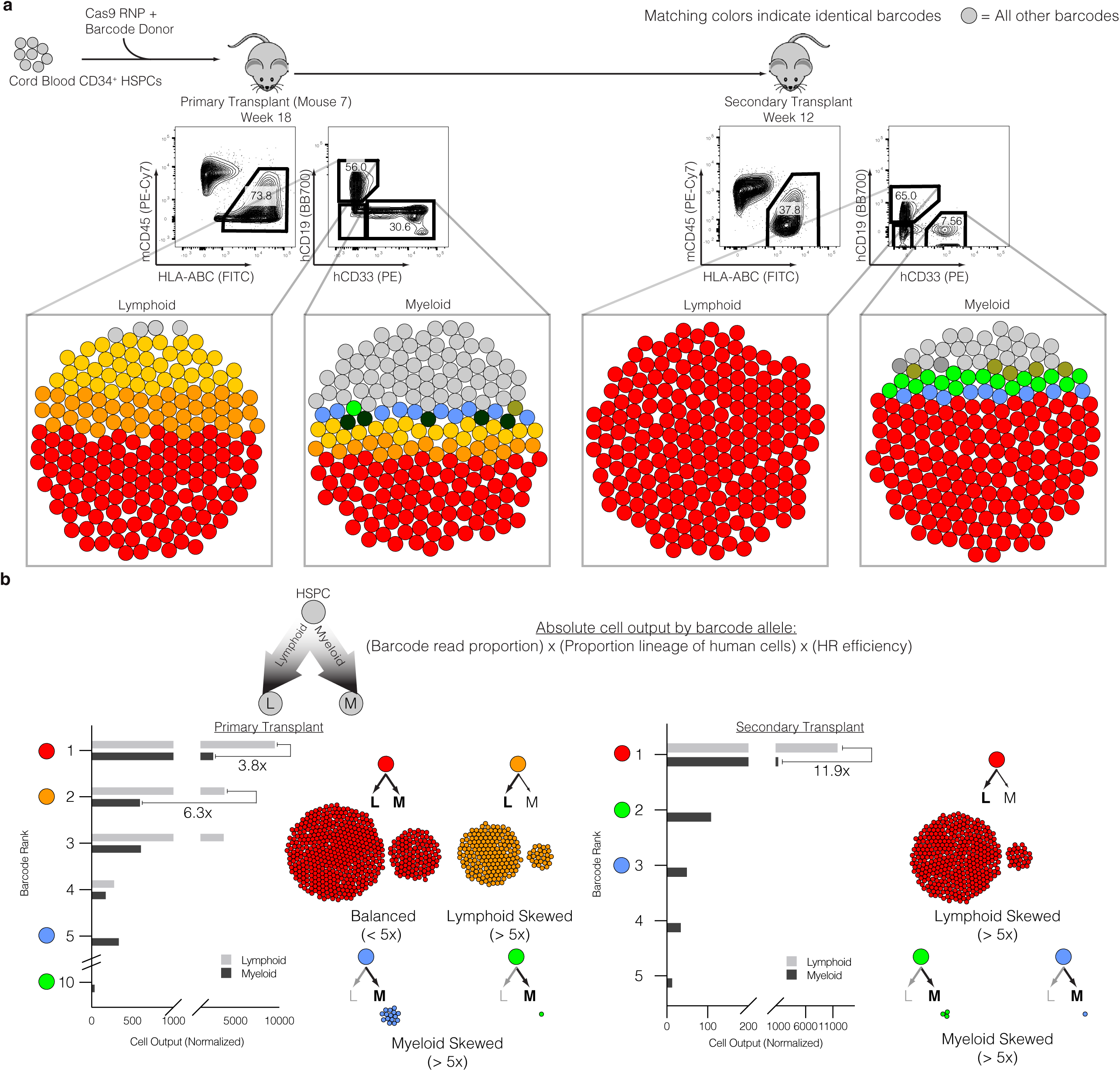
Clonal tracking of AAVS1 barcoded targeted HSPCs in reconstituting primary and secondary NSG transplants. **a** Top: Experimental schematic. Middle: Flow cytometry plots representing bi-lineage engraftment in primary transplant (left, week 18 post-transplant) and secondary transplant (right, week 12 post-transplant). Bottom: Bubble plots representing barcode alleles as unique colors from each indicated sorted population. Shown are the three most abundant clones from all six populations. All other barcodes represented as grey bubbles. **b** Normalized output of barcode alleles with respect to lineage contribution. Total cell output (bar graphs) from indicated barcodes adjusted for both differential lineage output and genome editing efficiency within each subset. Examples of various lineage skewing depicted, with cell counts reflecting relative contributions to the xenograft. One highly engrafted mouse (Mouse 7) depicted of n=5 total. Skewed output defined as 5-fold or greater bias in absolute cell counts towards lymphoid or myeloid lineages.

## Discussion

TRACE-Seq improves the understanding of the clonal dynamics of hematopoietic stem and progenitor cells following homologous recombination-based genome editing using two different gene targets (*HBB* and *AAVS1*). The data demonstrate that Cas9/sgRNA and AAV6 gene editing targets four distinct types of hematopoietic cells capable of engraftment, including: 1) rare and potent hematopoietic balanced LT-HSCs, 2) rare lymphoid skewed progenitors, 3) rare and potent myeloid skewed progenitors, and 4) more common and less proliferative myeloid skewed HSPCs.

TRACE-seq clearly demonstrates that in the NSG mouse model, engraftment of human cells after genome editing is largely oligoclonal with a few clones contributing to the bulk of hematopoiesis. From a technical perspective, we have developed a data analysis pipeline with multiple filters to distinguish sequencing artifacts from low abundance clones. As sequencing technologies and barcode design improve, the ability to distinguish noise from low abundance clones will similarly improve. Nonetheless, the evidence that clones that were seemingly rare in primary transplants can contribute significantly to hematopoiesis in secondary transplants demonstrates both the sensitivity of this method to detect such clones and the biologic importance of such clones in hematopoiesis.

We compare and contrast these results to lentiviral based genetic engineering of HSPCs since clonal dynamics of genome edited cells has not been published previously. Previous studies tracking LV IS in NSG mice have suggested on the order of 10-200 total clones (without data regarding the relative contributions of different clones) persisting long-term (although at different frequencies in each of the two mice analyzed), with identification of lineage-skewed as well as multi-potent LT-HSCs^50^. Accordingly, TRACE-Seq identified >50 clones per mouse that were contributing to the entire hematopoiesis of gene targeted cells (**Figure 3e**), suggesting that genome edited human HSPCs engraft as efficiently as lentiviral engineered cells in the NSG xenogeneic model. Interestingly, we identified 1-3 clones capable of robust multi-lineage reconstitution in secondary transplants, suggesting between one in 6 × 10^4^ and 4.6 x 10^5^ input cells are gene targeted LT-HSCs (based on the numbers of cells transplanted). In a clinical trial for Wiskott-Aldrich syndrome (WAS), IS analysis showed the frequency of CD34^+^ HSPCs with steady-state long term lineage reconstitutions falls between 1 in 100,000 and 1 in a 1,000,000 (a few thousand clones out of the ∼80-200 million HSPCs transplanted)^4^. Further building on this clinical trial, recent reports have suggested that LV integrations occur in cells within the HSPC pool that have long-term lymphoid or myeloid lineage restrictions as well^5^. Taken together, our data suggest that the frequency of gene-targeting and LV gene addition are similar in potent long-term engrafting LT-HSCs.

TRACE-Seq also demonstrated genome edited clones that were heavily lineage skewed in both primary and secondary transplants. This finding demonstrates that the gold standard of HSC function, namely serial transplantation, may not always identify multi-potent HSCs. Nonetheless, the method should allow assessment of other mouse xenograft models of human hematopoietic transplantation in supporting lineage restricted and multi-lineage reconstitution of genome edited cells, including models that further maintain healthy and leukemic myeloid and innate immune system development^51-53^. In the future, this method, potentially combined with novel cell sorting schemes to resolve lineage preference within the CD34+ fraction^54^, should help determine whether cells that undergo gene targeting have a bias towards particular lineages which may help guide which human genetic diseases of the blood may be most amenable to gene targeting based approaches. For example, if gene targeting preferentially occurs in long-term myeloid progenitors, this would support its use in diseases that require long-term myeloid engraftment of gene targeted cells such as sickle cell disease, chronic granulomatous disease, or beta-thalassemia.

In addition to helping understand hematopoietic reconstitution of genome edited cells in pre-clinical models, TRACE-Seq could also be used to further investigate the wide variety of genome editing approaches and HSC culture conditions to determine if they change either the degree of polyclonality or the lineage restriction of clones following engraftment. The wide numbers of variables that are under active study include different genome editing reagents and methods (different nucleases and donor templates and the inhibition of certain pathways^55, 56^), differing culture conditions (e.g. cytokine variations^57^, small molecules^58, 59^, peptides^55, 56^, and 3-D hydrogel scaffolds^60^), and altering the metabolic or cell cycle properties of the gene edited cells. This study, in which two different approaches targeting two different genes was established, serves as the key foundation for such future studies.

In conclusion, TRACE-seq demonstrates that homologous recombination-based genome editing can occur in human hematopoietic stem cells as defined by multi-lineage reconstitution following serial transplantation at a single cell, clonal level. Moreover, TRACE-Seq lays the foundation of clonal tracking of gene targeted HSPCs for basic research into normal and malignant hematopoiesis. The ability of track clones in a clinical setting has proven to be a powerful approach to understand the safety, efficacy, and clonal dynamics of lentiviral based gene therapies, and it will be informative to determine if regulatory agencies will accept having innocuous barcodes as part of recombination donor templates in clinical studies so that the safety, efficacy, and clonal dynamics of reconstituted gene targeted cells, including HSCs, T-cells, or other engineered cell types, can be tracked following administration to patients.

## Figure Legends

**Extended Data Figure 1.**
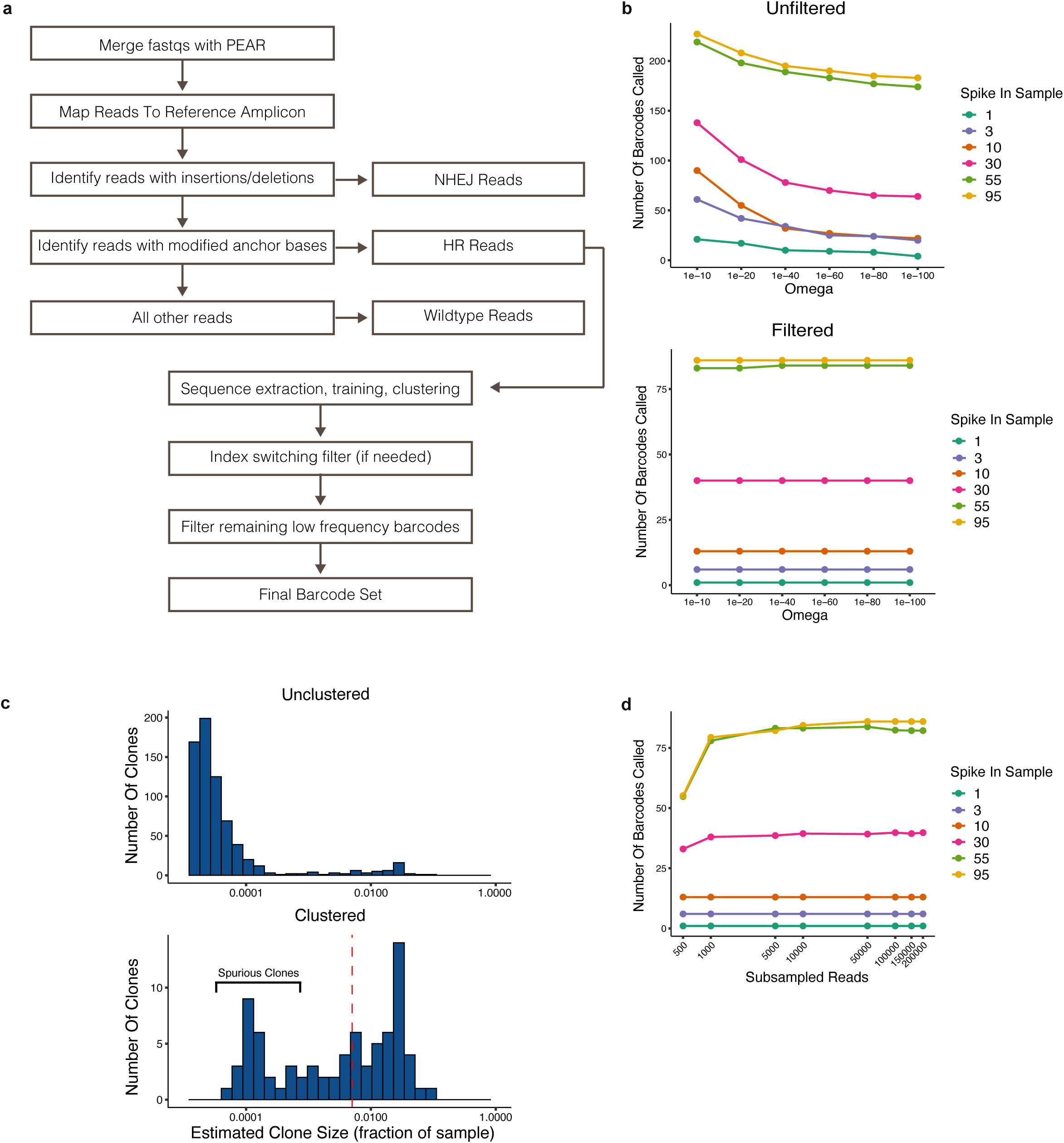
TRACE-Seq barcode analysis pipeline and optimization. **a** Schematic of barcode calling pipeline workflow. **b** Number of barcodes called from sequencing of spike-in samples (**Figure 1e**) as a function of the DADA2 Omega parameter before filtering barcodes <0.5% (top) and after filtering barcodes (bottom). **c** Distribution of barcode sizes for a sample with a known barcode content of 30 barcodes. Top: distribution of unclustered barcodes, showing large numbers of very infrequent sequences on the left side of the distribution. Bottom: distribution of clustered barcodes (bottom). Threshold of 0.5% is depicted by the dashed red line, which was used to enrich for “high confidence” barcodes in subsequent analyses. **d** Subsampling analysis of spike-in samples to estimate necessary sequencing depths to recover the expected numbers of barcodes.

**Extended Data Figure 2.**
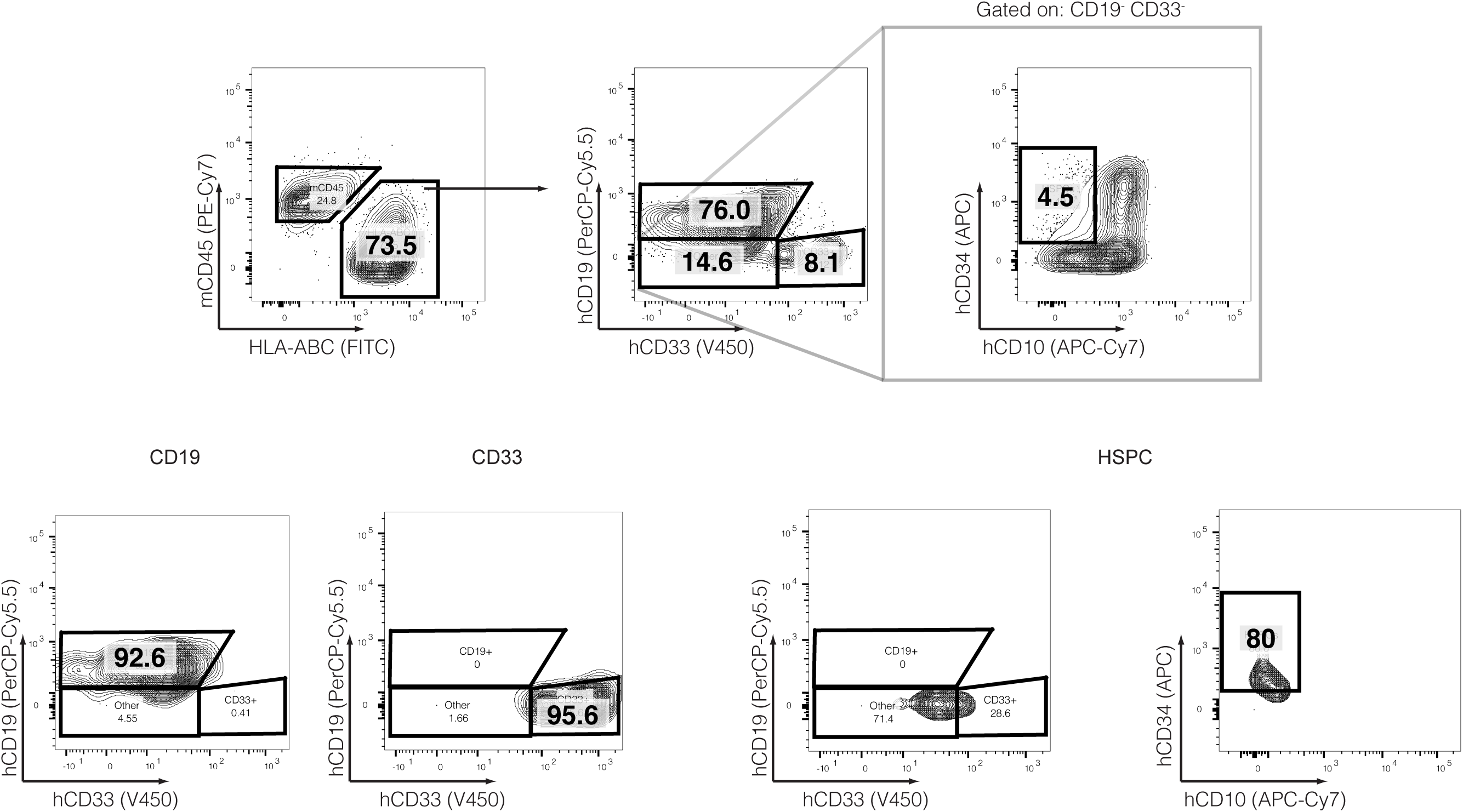
Representative HBB gating strategy (upper) and sort purity (lower) for CD19^+^, CD33^+^, and HSPCs (CD19^-^CD33^-^CD10^−^CD34^+^) populations.

**Extended Data Figure 3.**
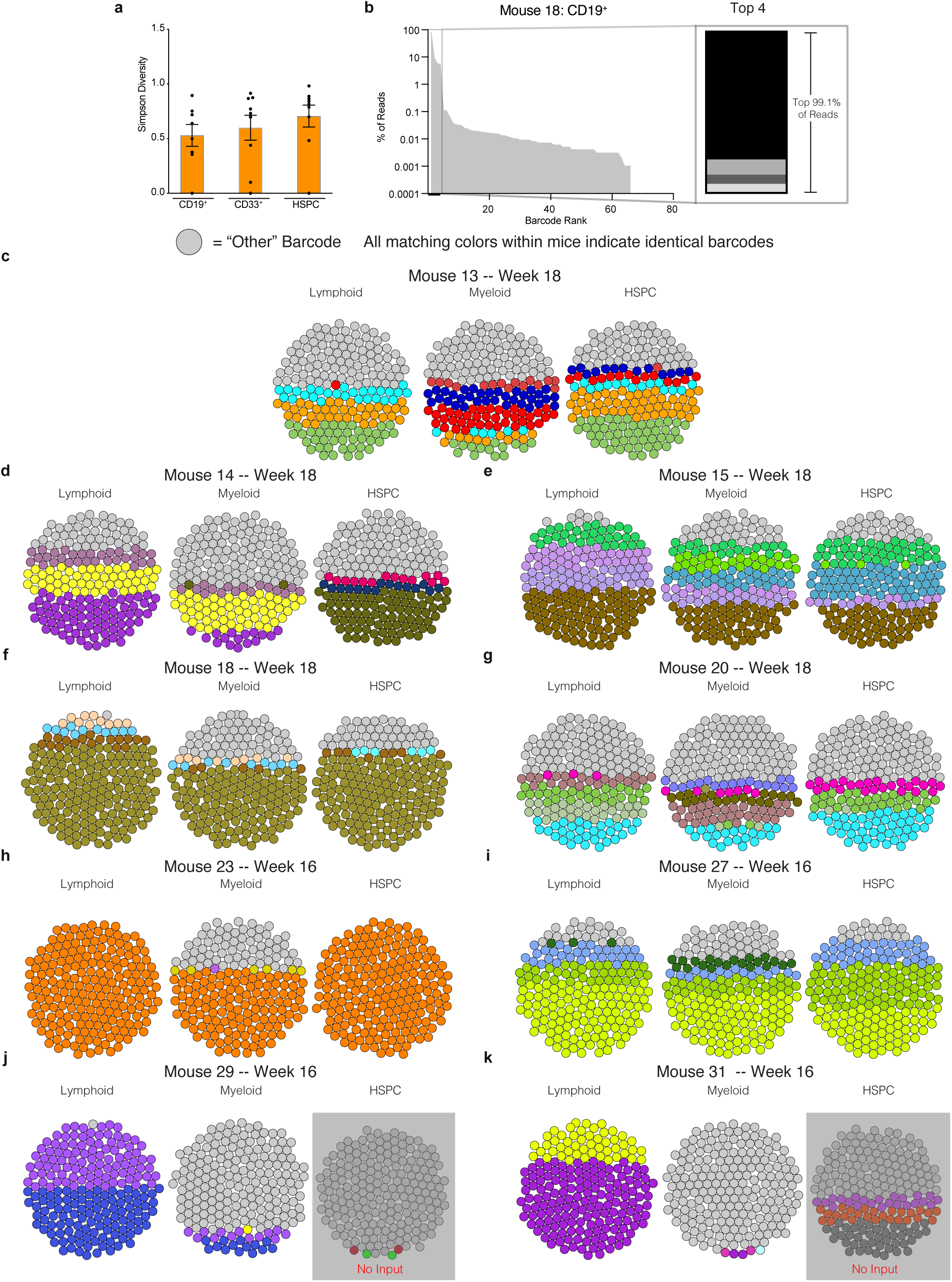
Bubble plots depicting shared and unique barcode representation in all HBB barcoded mice at sacrifice. **a** Simpson diversity, representing barcode richness and evenness, for each lineage. **b** Left: representative quantification of all barcodes in descending order by read proportion. Right: Top four barcodes depicted as stacked bar chart. **c-k** Visualization of top 3 barcodes from all sorted populations (similar to **Figure 4a**) from indicated mice. All other barcodes represented as grey bubbles. Error bars depict mean ± SEM. Note: Despite some colors appearing similar, no top barcodes are shared between distinct mice. See **Supplemental Table 3** for individual HBB sequences and barcode counts.

**Extended Data Figure 4.**
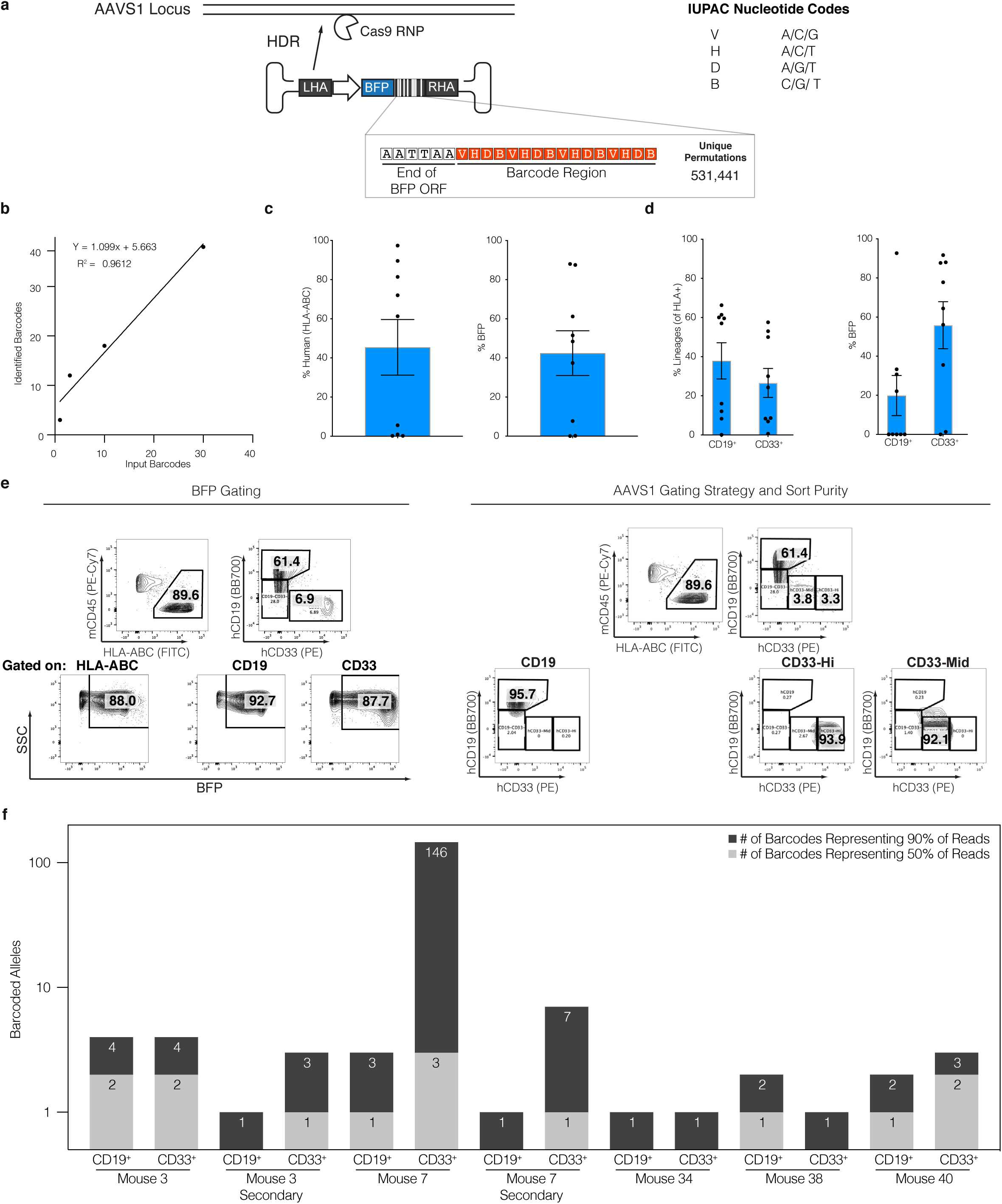
Engraftment clonality with genome edited HSPCs is independent of locus and barcode diversity. **a** Top: Schema of AAVS1 locus and AAV6 donor containing SFFV-BFP-[barcode]-pA expression cassette. Bottom: Barcode region depicted in red, with a maximum theoretical diversity of 3^12^ = 531,441. **b** Recovery of barcodes from untreated genomic DNA containing 1, 3, 10, and 30 individual plasmids containing BFP barcodes. **c** Total human engraftment in whole bone marrow collected 16-18 weeks post transplantation (expressed as proportion of human HLA-ABC^+^ cells), left, and genome editing efficiency as measured by percentage of BFP+ cells by flow cytometry, right. **d** Multi-lineage engraftment of human CD19^+^ and CD33^+^ lineages, left, with respective genome editing efficiencies, right. Error bars depict mean ± SEM. **e** Example of gating on highly engrafted mouse on %BFP^+^ within each lineage (left) and representative gating strategy and sort purity (right). **f** Barcodes from each subset were sorted from largest to smallest by percentage of reads. Depicted are the numbers of most abundant, unique barcode alleles comprising the top 50% and top 90% of reads from each lineage of all mice transplanted with BC donor edited HSPCs. Mean ± SEM genomes analyzed from each group— CD19^+^: 7900 ± 1500, CD33^+^: 10000 ± 3000 (see **Supplemental Table 2c**).

**Extended Data Figure 5.**
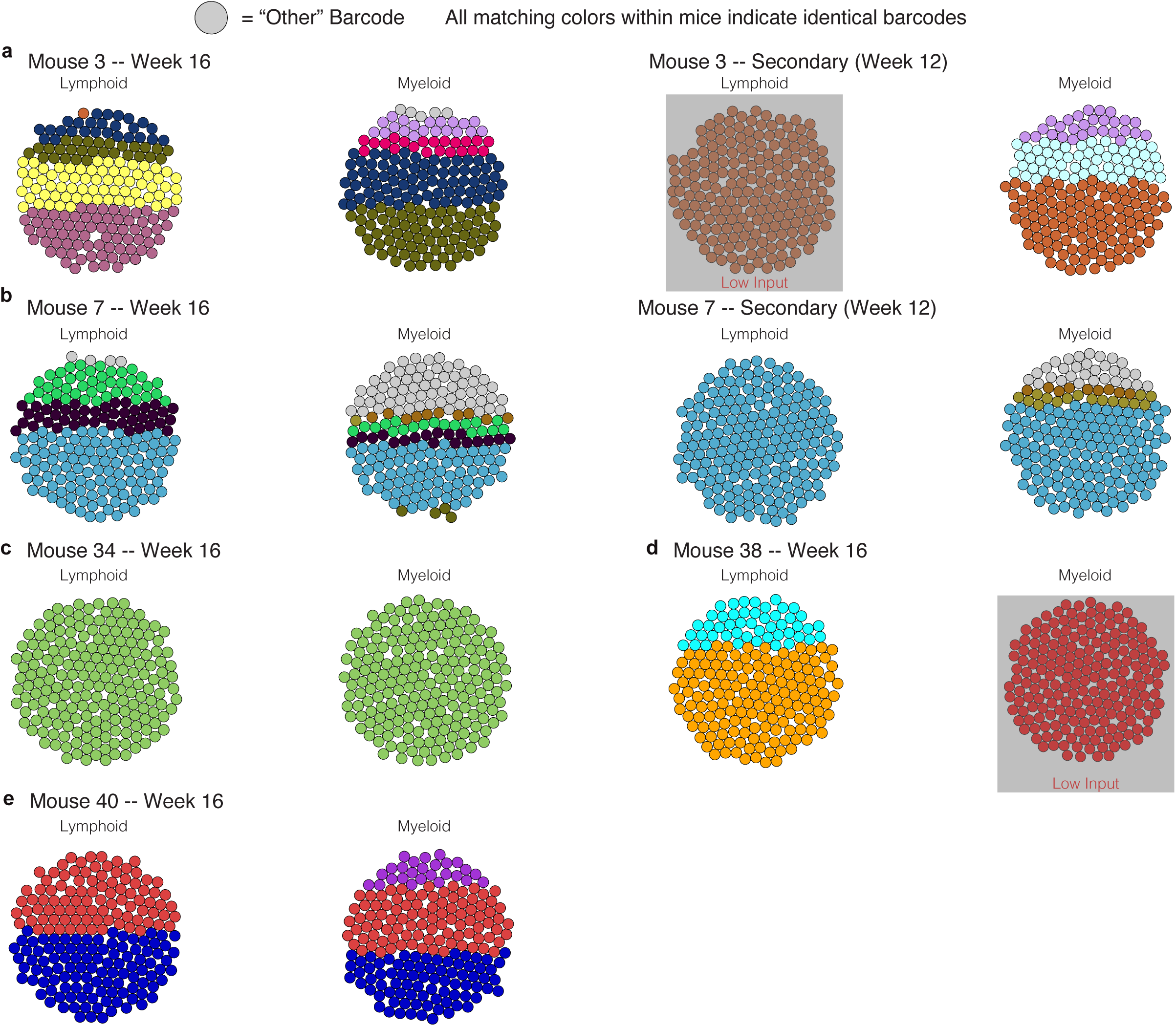
Bubble plots depicting shared and unique barcode representation in all BFP barcoded mice at sacrifice. **a-e** Visualization of top 3 barcodes from sorted populations (similar to **Figure 5a**) from indicated mice. All other barcodes represented as grey bubbles. Despite some colors appearing similar, no top barcodes are shared between distinct mice. See **Supplemental Table 4** for individual BFP sequences and barcode counts.

**Supplemental Table 1.**
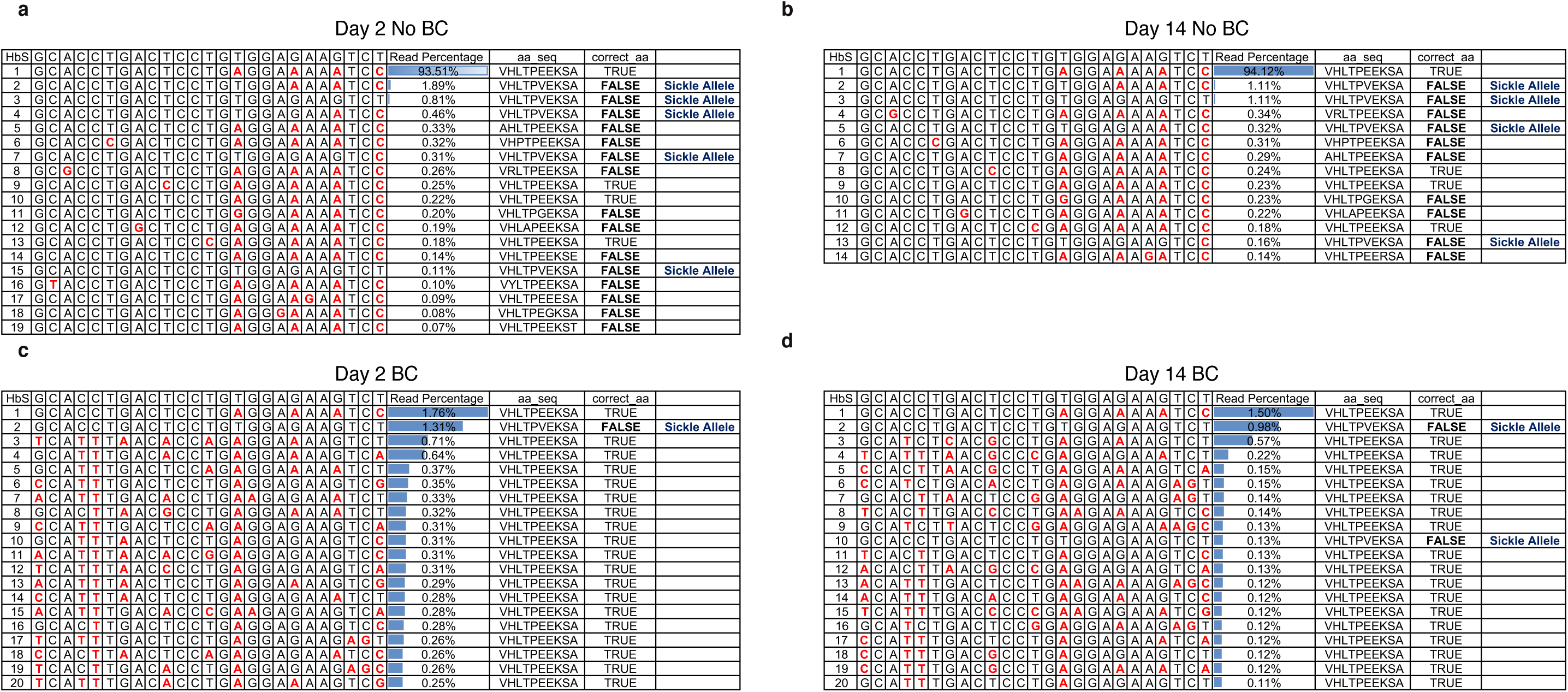
HBB barcodes identified in vitro, related to **Figure 2c**. Barcodes colorized with red nucleotides indicating differences from unmodified sequence (HbS, top row), along with read percentages, amino acid translation (“aa_seq”), and a True/False value indicating whether the HBB protein translation matches the WT sequence (“correct_aa”).

**Supplemental Table 2.**
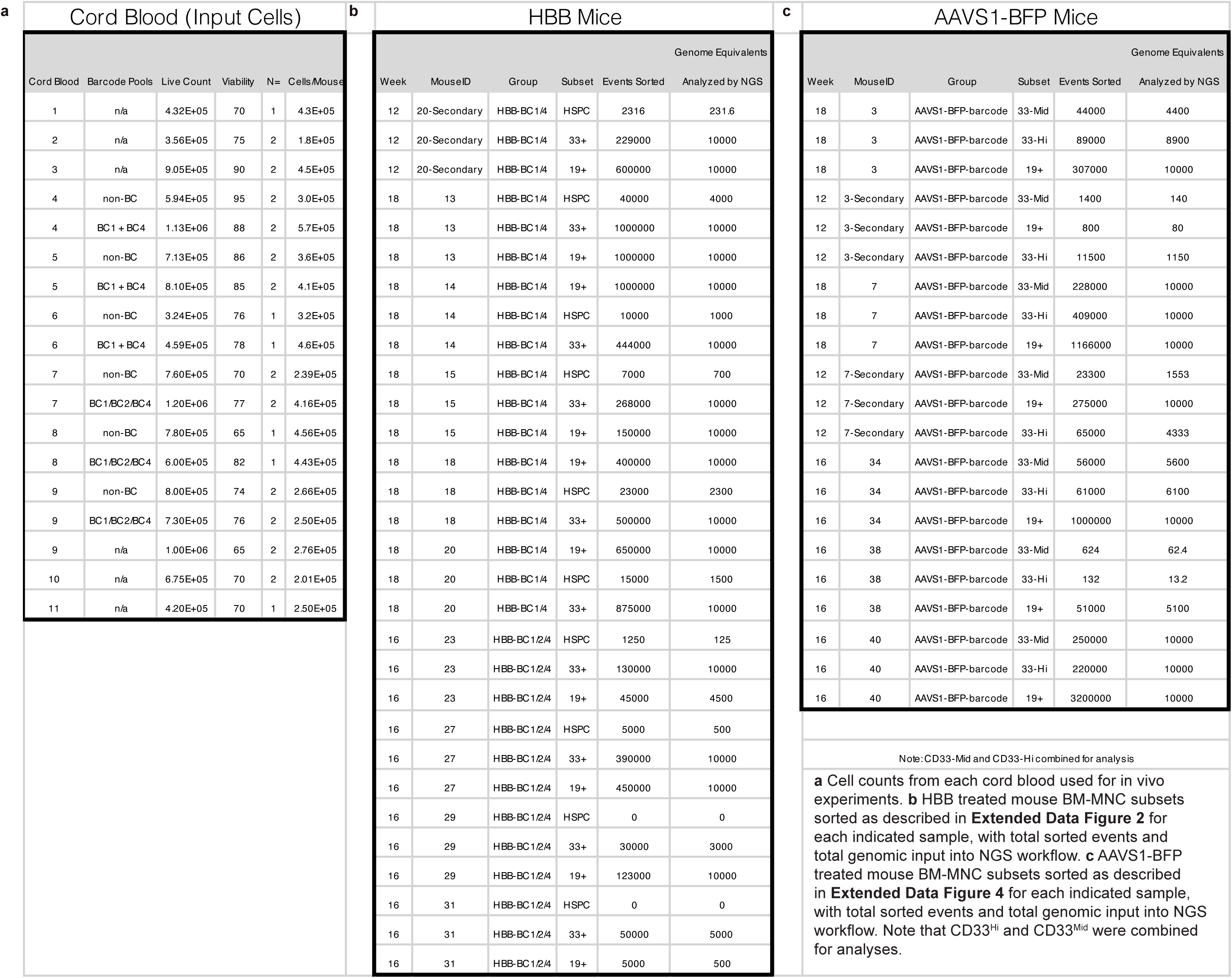
In Vivo Cell Transplants and Sort Data

**Supplemental Table 3.**
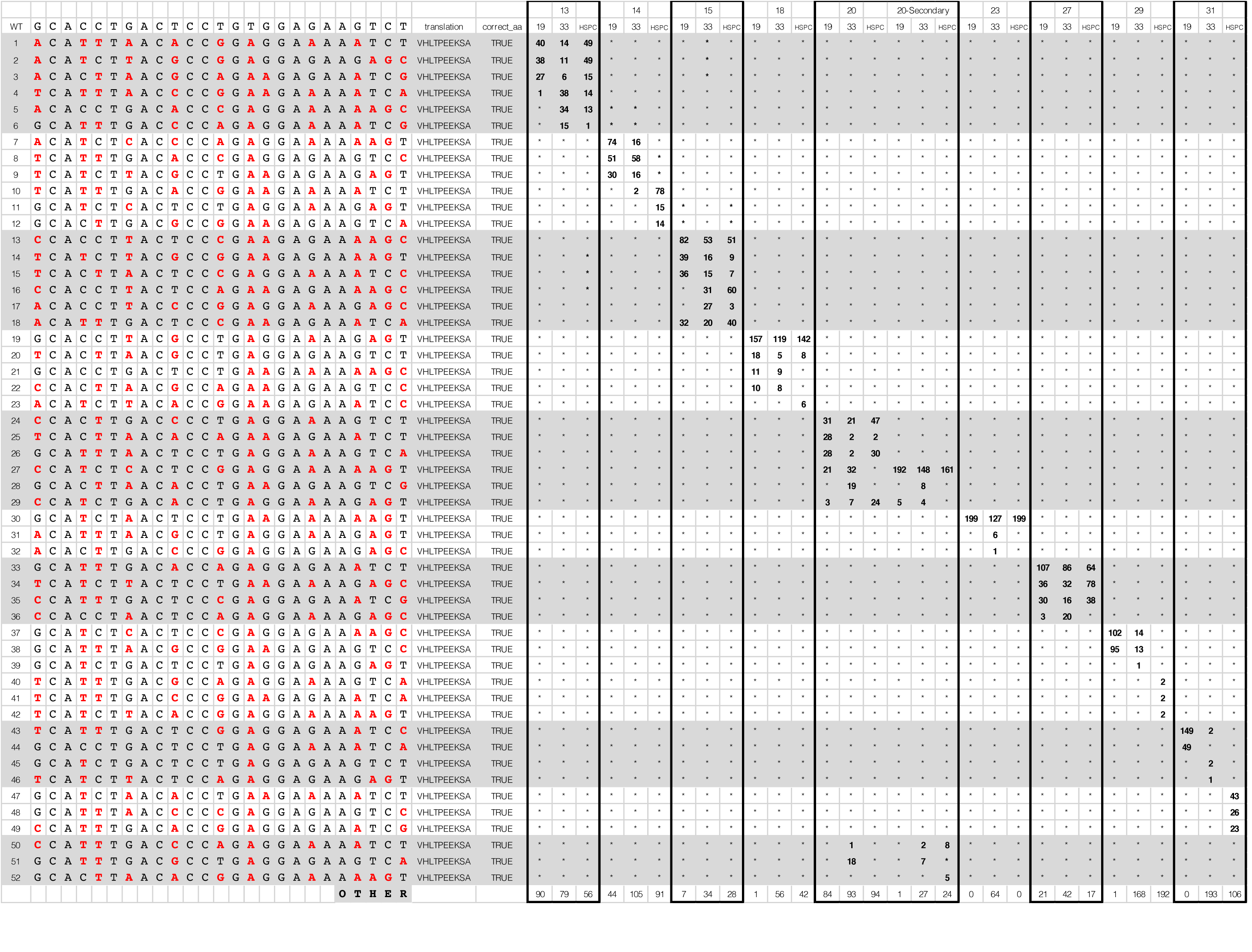
HBB “Bubble plot” detailed legend. Total counts and barcode identities for barcodes depicted in **Figure 4** and **Extended Data Figure 3**.

**Supplemental Table 4.**
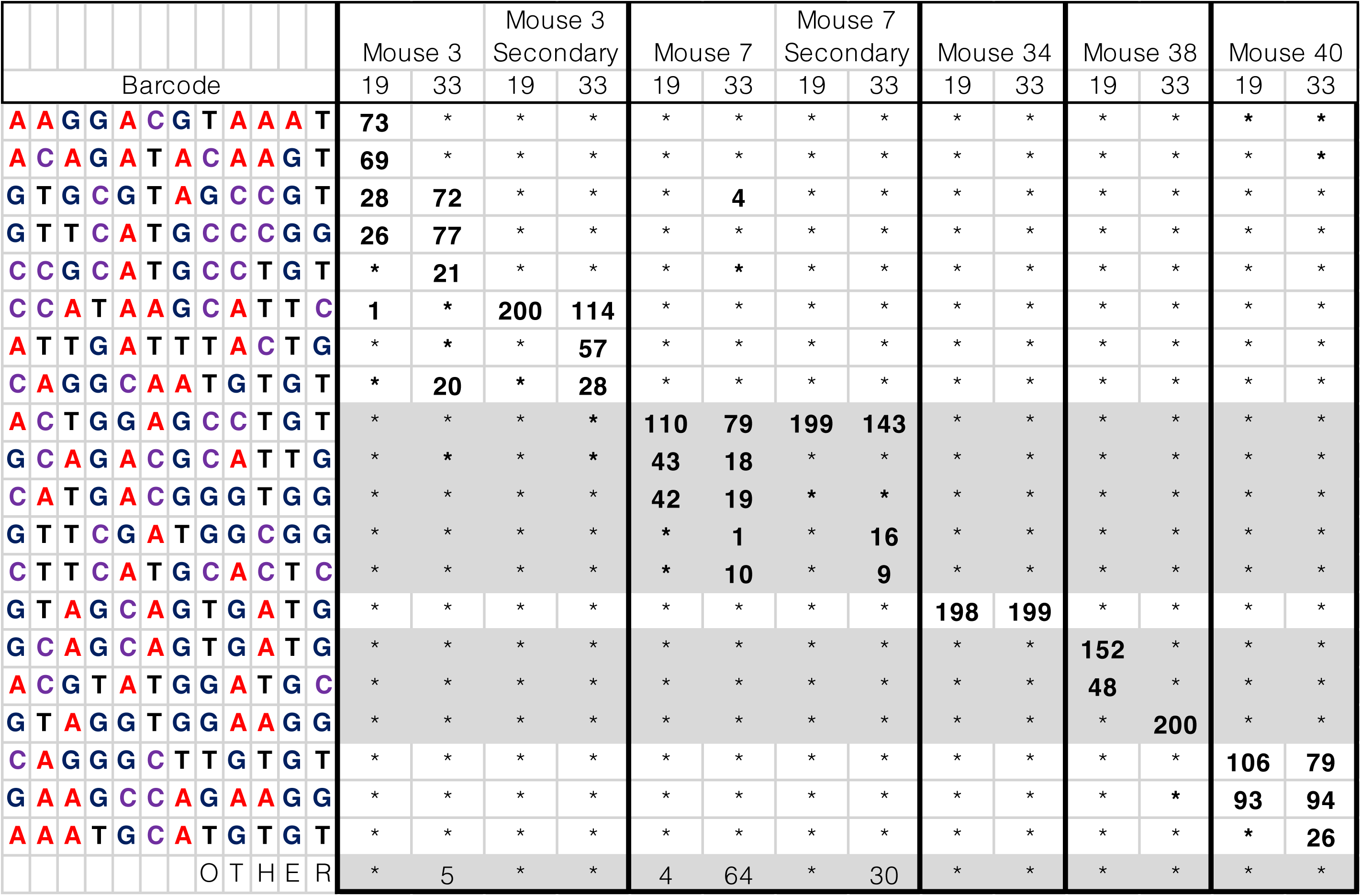
BFP “Bubble plot” detailed legend. Total counts and barcode identities for barcodes depicted in **Figure 5** and **Extended Data Figure 5**.

**Supplemental Video 1**

Screen recording of R Shiny app for generating and interacting with bubble plot data as in **Figures 4 and 5** and **Extended Data Figures 3 and 5**.

## Acknowledgements

R.S. was supported by a Postdoctoral Fellowship, PF-18-102-01-DOC, from the American Cancer Society. R.M. acknowledges support for the Stanford Ludwig Center for Cancer Stem Cell Research and Medicine, and NIH grants R01-CA188055 and R01-HL142637. M.H.P. gratefully acknowledges the support of the Amon Carter Foundation, the Laurie Kraus Lacob Faculty Scholar Award in Pediatric Translational Research and NIH grant support R01-AI097320 and R01-AI120766. G.B. acknowledges support from the Cancer Prevention and Research Institute of Texas (RR14008 and RP170721). D.P.D and J.C. were supported by NIH grant support R01-HL135607. We thank the Binns Program for Cord Blood Research at Stanford University for cord-blood-derived CD34^+^ HSPCs and also for SCD-HSPCs. Patients with SCD consented to the use of CD34^+^ HSPCs for research with the accompanying IRB approval. All mouse experiments were conducted in accordance with a protocol approved by the Institutional Animal Care and Use Committee (Stanford Administrative Panel on Laboratory Animal Care no. 22264).

## Author Contributions

R.S., D.P.D., R.M., and M.H.P. conceived the study. R.S. and D.P.D. designed and performed experiments. A.A. designed and performed bioinformatic analyses. C.M.L., and Y.P. performed the library preparation for MiSeq and HiSeq runs. J.C. and T.K. developed reagents and performed analyses. R.S., D.P.D., A.A., R.M., and M.H.P. wrote the manuscript with support from all authors.

## Competing Financial Interests

M.H.P. has equity and serves on the scientific advisory board of CRISPR Therapeutics and Allogene Therapeutics. R.M. is a founder, consultant, equity holder, and serves on the Board of Directors of Forty Seven Inc. However, none of these companies had input into the design, execution, interpretation, or publication of the work in this manuscript.

## Data and code availability

Barcode counts after processing by the TRACEseq pipeline will be made available on the GEO database. Code used to generate bubble plots will be available on github (https://github.com/armonazizi/TRACEseq).

